# Region-specific Nucleus Accumbens Dopamine Signals Encode Distinct Aspects of Avoidance Learning

**DOI:** 10.1101/2024.08.28.610149

**Authors:** Gabriela C. Lopez, Louis D. Van Camp, Ryan F. Kovaleski, Michael D. Schaid, Oscar Andrés Moreno-Ramos, Rajeshwar Awatramani, Julia M. Cox, Talia N. Lerner

## Abstract

Avoidance learning – learning to avoid bad outcomes – is an essential survival behavior. Dopamine signals are widely observed in response to aversive stimuli, indicating they could play a role in learning about how to avoid these stimuli. However, it is unclear what computations dopamine signals perform to support avoidance learning. Furthermore, substantial heterogeneity in dopamine responses to aversive stimuli has been observed across nucleus accumbens (NAc) subregions. To understand how heterogeneous dopamine responses to aversive stimuli contribute to avoidance learning, we recorded NAc core (Core) and NAc ventromedial shell (vmShell) dopamine during a task in which mice could avoid a footshock punishment by moving to the opposite side of a 2-chamber apparatus during a five-second warning cue. We found that NAc Core and vmShell dopamine signals responded oppositely during shocks and warning cues. Both signals evolved substantially – but differently – with learning. NAc vmShell dopamine responses to cues and shocks were present during early learning but not sustained during expert performance. NAc Core dopamine responses strengthen with learning and are especially evident during expert performance. Our data support a model in which NAc vmShell dopamine guides initial cue-shock associations by signaling salience, while NAc Core dopamine encodes prediction errors that guide the consolidation of avoidance learning.

## Introduction

Avoidance learning is essential for survival. Whether it is a fire alarm warning of a nearby fire, or the rustle of leaves warning of a nearby predator, animals must learn to heed cues that predict danger and act to preserve their wellbeing. When performed in excess, however, avoidance can become maladaptive, such as in anxiety disorders, obsessive-compulsive disorder (OCD), and depression.^1,2^ To better understand how avoidance learning contributes to these conditions, it is important to understand the underlying neural mechanisms.

Avoidance learning is a form of instrumental learning, in which animals learn from negative reinforcement (the removal or avoidance of something bad). Dopamine is well known to be essential for instrumental learning, but it is best studied in the context of positive reinforcement (the receipt of something good).^3,4^ However, dopamine neurons respond not only to rewarding but also to aversive stimuli and associated cues, raising the possibility that they play an important role in learning aversive associations to facilitate the performance of avoidance actions.

Dopamine neurons respond heterogeneously to aversive stimuli,^5–9^ leading to debate about the computations being performed. Increases in dopamine in response to aversive stimuli have been proposed to reflect salience encoding whereas decreases are largely considered consistent with valence or reward prediction error encoding.^10–14^ Both types of information might support avoidance learning, but little evidence is available as most studies of dopamine signaling in response to aversive stimuli utilize paradigms with inescapable exposure to aversive stimuli and do not allow avoidance. Evidence that is available supports a role for dopamine in avoidance learning,^15–19^ but the region specificity and evolution of dopamine responses during learning remain in question.^20^

Region specificity is an important parameter to consider when studying dopaminergic signaling. A previous study that carefully mapped the region specificity of dopaminergic responses to aversive stimuli in the nucleus accumbens (NAc) during Pavlovian learning found that dopamine release in the NAc ventromedial shell (vmShell) is increased, where dopamine release in most other NAc subregions (including the Core, dorsomedial Shell and lateral Shell) is decreased.^7,21^ NAc vmShell is heavily innervated by the amygdala, hypothalamus, and hippocampus, structures that are important for fear and emotional processing,^22–25^ and its function is often associated with the integration of emotional and motivational value.^10,26,27^ In contrast, the NAc Core is important for attention and contextual learning.^28–30^ NAc Core dopamine has been hypothesized to encode prediction errors^13,31,32^ and may specifically encode safety prediction errors during avoidance learning.^16^

Here, we asked: how do NAc vmShell and Core dopamine signals evolve as animals learn to reduce or prevent an aversive outcome? How do these signals differ when the occurrence of punishment is controllable vs uncontrollable? We found that NAc vmShell and Core dopamine signals evolve across avoidance learning, with distinct changes observed in early and expert stages, respectively. Our data shed new light on how the dopamine system supports the acquisition of avoidance behavior and offer new hypotheses regarding the computational roles played by distinct dopaminergic circuits during instrumental learning from aversive stimuli.

## Methods

### Mice

Adult female and male mice were group-housed by sex under a conventional 12:12 h light/dark cycle and were given ad libitum access to food and water. Wildtype C57BL6/J mice (Jackson Strain #:000664) were used for vmShell experiments. Heterozygous Tg(Adora2a-Cre)KG139Gsat and Tg(Drd1-Cre)FK150Gsat mice, generated by crossing homozygous Tg(Adora2a-cre)KG139Gsat and Tg(Drd1-Cre)FK150Gsat mice with WT (C57BL6/J) in-house, were used for Core experiments. To distinguish Sox6+ and Sox6-dopamine axons, DAT-2A-Flpo (unpublished, generated by Dr. Awatramani) or TH-2A-Flpo^33^ mice were crossed with Sox6-FSF-Cre^34^ mice. Mice heterozygous for DAT-2A-Flpo or TH-2A-Flpo and Sox6-FSF-Cre were used for experiments. Littermates were randomly assigned to experimental groups (Core: 5 females, 9 males; vmShell: 5 females, 3 males; Sox6: 3 females, 4 males). All experiments were approved by the Northwestern University Institutional Animal Care and Use Committee.

### Surgery

Viral injection and optic fiber implant surgeries were performed on adult mice at 7-10 weeks of age. Mice were anesthetized in an isoflurane chamber at 3-4% isoflurane (Henry Schein) and then placed on a stereotaxic frame (Stoelting). Anesthesia was maintained at 1-3% isoflurane. Mice were injected with meloxicam (Covetrus, 20 mg/kg) subcutaneously prior to the start of surgery to minimize post-surgical pain. Hair was removed from the top of the head using Nair. The exposed skin was disinfected with alcohol and a povidone-iodine solution. Prior to incision, bupivacaine (Hospira, 2 mg/kg) was injected subcutaneously at the incision site. The scalp was opened using a sterile scalpel and holes were drilled in the skull at the appropriate stereotaxic coordinates. Viruses were infused at 100 nL/min through a blunt 33-gauge injection needle using a syringe pump (World Precision Instruments). The needle was left in place for 5 min following the end of the injection, then slowly retracted to avoid leakage up the injection tract. Implants were secured to the skull with Metabond (Parkell) and Flow-it ALC blue light-curing dental epoxy (Pentron). After surgery, mice were allowed to recover until ambulatory on a heated pad, then returned to their homecage with moistened chow or DietGel available. The mice were checked after 24 hours and provided with another dose of meloxicam. Mice then recovered for three to four weeks before behavioral experiments began.

A2A-Cre and D1-Cre mice were injected with a 500 nL virus cocktail containing AAV9-CAG-dLight1.3b (1.25e12 vg/ml, Virovek) and AAV1-CAG-FLEX-NES-jRCaMP1b (2.6e12 vg/ml, Addgene) into the Core (AP 1.5, ML 0.9, DV -4.1) in one hemisphere. Hemispheres were counterbalanced between mice. Fiber optic implants (Doric Lenses; 400 µm, 0.48 NA, 1.25 mm ferrules) were placed above the Core (AP 1.5 mm, ML 0.9 mm, DV -4.0 mm) in the same injection hemisphere. WT mice were injected with 500 nL of AAV9-CAG-dLight1.3b (1.25e12 vg/ml, Virovek) into the vmShell (AP 1.5, ML 0.6, DV -4.8) in one hemisphere. Hemispheres were counterbalanced between mice. Fiber optic implants (Doric Lenses; 200 µm, 0.48 NA, 1.25 mm ferrules) were placed above the vmShell (AP 1.5 mm, ML 0.6 mm, DV -4.7 mm) in the same injection hemisphere.

For fiber photometry experiments, Dat-2A-Flpo x Sox6-FSF-Cre and TH-2A-Flpo x Sox6-FSF-Cre mice were injected with 500 nL of AAV-Ef1α-Con/Fon-GCaMP6f (2e13 vg/ml, Addgene) or AAV-Ef1a-Coff/Fon-GCaMP6f (1.9e13 vg/ml, Addgene) into the Core (AP 1.5 mm, ML 0.9 mm, DV -4.1 mm) in one hemisphere to express GCaMP in Sox6+ or Sox6-expressing VTA dopamine terminals, respectively. Hemispheres were counterbalanced between mice. Fiber optic implants (Doric Lenses; 400 µm, 0.48 NA, 1.25 mm ferrules) were placed above the Core (AP 1.5, ML 0.9, DV -4.0) in the same injection hemisphere. Dopamine release dynamics and calcium signals from ventral tegmental area (VTA) terminals in the NAc were recorded for all 8 days of behavior.

For histology only (Figure 8A), Sox6-FSF-Cre/Dat-2A-Flpo adult mice (P>60) were anesthetized with 3% isoflurane and placed onto a stereotaxic apparatus. They were kept under anesthesia with ∼1.5% isoflurane. A volume of 0.45uL of virus was slowly injected over 5 minutes into the VTA (AP -3.16mm, ML : -0.45mm, DV: -4.10mm) using a Hamilton Neuros syringe filled with a viral cocktail mixture (1:1) of AAV8 HSyn-Con/Fon-YFP (2.4e13 vg/mL, Addgene) and AAV8 Ef1a-Coff/Fon-mCherry (2.2e13 vg/mL, Addgene). The needle was removed 5 minutes after the injection. The cranial wound was closed with staples. Animals were perfused 4 weeks after the injection.

### Behavior

Mice underwent two shock-paired behavioral paradigms: active avoidance and inescapable shock. Both tests were completed using custom 2-chamber shuttle boxes, with chambers separated by a plastic door (MED Associates). The day before testing began, mice were habituated to tethering with patch cords (Doric Lenses). During active avoidance, mice were tethered with patch cords and then placed in either the right or left shuttle box chamber. Once a mouse was detected in a chamber, a sound and light cue turned on for 5 s. If the mouse shuttled from its initial chamber to the opposite chamber within 5 s, the light and sound cue turned off. If the mouse failed to cross over to the opposite chamber within 5 s, a 0.4 mA shock turned on and continued for 25 s or until the mouse shuttled to the opposite chamber. Shuttling after shock turned off the sound and light cue as well. The task resulted in two major behavioral measures: 1) Avoided Shocks – in which the mouse moved to the opposite chamber before the shock began (<5 s after cue start), and 2) Escaped Shock – in which the mouse moved to the opposite chamber after the shock had begun (>5 s after cue start). Mice were tested on this task for 7 days, 30 trials per day. There was an inter-trial interval (ITI) between every trial that lasted an average of 45 s. After active avoidance testing, mice underwent one day of inescapable shock. For this task, mice were tethered and placed within the 2-chamber shuttle box as previously described. Once a mouse was detected in a chamber, a sound and light cue turned on for 5 s. After 5 s, a 0.4 mA shock turned on and remained on for 5 s. Mice were able to shuttle between chambers, but this behavior did not prevent or stop the shock. After 5 s, the shock and cues turned off. Mice were tested on this task for 1 session consisting of 10 trials.

### Video Analysis

Mouse pose estimation was conducted using SLEAP.^35^ Body part points (nose, left ear, right ear, mid-body, left body, right body, and tail base) were manually placed on sample video frames to train a model to accurately predict body part locations. One to two hundred labels were added at a time to generate intermediate models and assess accuracy. Ultimately, 679 frames across all videos were labeled to achieve human-like labeling accuracy with minimal artifacts. The top 1% of predicted movements (corresponding to those > 480cm/s) were removed as artifacts largely stemming from frames where the mouse was not detected by SLEAP. Nose points generated from the SLEAP model were used for behavioral classification of freezing using Simple Behavior analysis (SimBA) software.^36^ Velocity of the nose was used to determine freezing, defined as <0.22cm/frame (6.6cm/s) movements for at least 2s in R.

### Fiber photometry

All recordings were performed using a fiber photometry rig with optical components from Doric lenses and Tucker Davis Technologies (TDT) controlled by a real-time processor from TDT (RZ5P or RZ10X). TDT Synapse software was used for data acquisition. 465nm, 405nm, and 560nm LEDs were modulated at 210 Hz, 330 Hz, and 450 Hz, respectively, for Core A2A-and D1-Cre probes. 465nm and 405nm LEDs were modulated at 210 Hz and 330 Hz, respectively for vmShell and Core Sox6+/-experiments. LED currents were adjusted to return a voltage between 150-200mV for each signal, were offset by 5 mA, were demodulated using a 4 Hz lowpass frequency filter. Behavioral timestamps, e.g., for cue and shuttle crossings, were fed into the real-time processor as TTL signals from the shuttle boxes (MED Associates) for alignment with the neural data. GuPPy, an opensource Python-based photometry data analysis pipeline, was used to determine dLight1.3b transients and GCaMP6f activity aligned to specific time-locked events.^37^

### Perfusions and histology

Mice received lethal i.p. injections of Euthasol (Virbac, 1mg/kg) to induce a rapid onset of unconsciousness and death. Once unresponsive to a firm toe pinch, an incision was made up the middle of the body cavity. An injection needle was inserted into the left ventricle of the heart, the right atrium was punctured, and solution (PBS followed by 4% PFA) was infused into the left ventricle as the mouse was exsanguinated. The mouse was then decapitated, and its brain was removed and fixed overnight at 4°C in 4% PFA. After perfusion and fixation, brains were transferred to a solution of 30% sucrose in PBS (w/v), where they were stored for at least two overnights at 4°C before sectioning. Tissue was sectioned on a freezing microtome (Leica) at 30 µm, stored in cryoprotectant (30% sucrose, 30% ethylene glycol, 1% polyvinyl pyrrolidone in PB) at 4°C until immunostaining. Dat-2A-Flpo x Sox6-FSF-Cre adult mice (P>60) were perfused with cold 4% PFA in PBS pH 7.4, the brains were dissected and fixed overnight in the same solution, cryoprotected in 30% sucrose in PBS, sectioned at 25 μm using a freezing microtome, and stored at –20°C in cryoprotectant solution.

Anti-GFP staining was performed on free floating sections to amplify signals from dLight1.3b and GCaMP6f. Sections were blocked in 3% normal goat serum in PBS for 1 h at room temperature. Primary antibody staining was performed using 1:500 Rabbit anti-GFP primary antibody (Invitrogen, A11122) in blocking solution at 4°C for 48 hrs. Secondary staining was performed using 1:500 donkey anti-rabbit Alexa Fluor 647 (Jackson ImmunoResearch, 711-606-152) in blocking solution at room temperature for 2 hrs. Tissue was mounted on slides in PBS and coverslips (Fisherbrand, Cat. No. 1255005) mounted with DAPI Fluoromont-G (Southern Biotech). Slides were imaged using a fluorescent microscope (Keyence BZ-X710) with 5x, 10x, and 40x air immersion objectives. Probe placements were determined by comparing their location to the Allen Mouse Brain Atlas.

### Quantification and Statistics

#### Behavioral analysis

Behavioral data such as number of avoided or escaped trials and latencies to cross were collected automatically by MED-PC software (Med Associates). Percent avoidance was calculated by taking the number of avoided trials and dividing it by the total number of trials in the task (30 trials), then multiplying by 100. Freezing and inter-trial interval (ITI) crossing data were extracted via SLEAP^35^ for animal pose tracking and SimBA^36^ for behavioral classification.

#### Fiber photometry analysis

GuPPy^37^ was used to analyze dLight1.3b and GCaMP6f signals time-locked to specific behavioral events using default settings. In brief, raw data were passed through a zero-phase digital filter and a least-squares linear fit was applied to the 405nm control signal to align it to the 465nm or 560nm signal. ΔF/F was calculated with the following formula: (signal -fitted control) / (fitted control). To facilitate comparisons across animals, z-scores were calculated by subtracting the mean ΔF/F calculated across the entire session and dividing by the standard deviation (GuPPy standard z-score method). Peristimulus time-histogram (PSTH) parameters were set from -25 to 20 seconds and baseline parameters were set to from -5 to -2 seconds. H5 output files from GuPPy were imported into MATLAB for further quantitative analysis. Area-under-the-curve (AUC) was calculated in MATLAB using the trapz function. Dip amplitudes were calculated in MATLAB by inverting the plots about the y-axis and using the max and findpeaks functions.

#### Encoding model of dopamine fluorescence

To relate dopamine activity to behavioral events while accounting for the linear contributions of other events occurring close in time, we built a kernel-based encoding model of the photometry signals. For each session, we first downsampled the photometry traces to 20 Hz and then concatenated the photometry data across mice. We then used a multiple linear regression analysis with the photometry signal as the dependent variable and behavioral events as independent variables. To account for lags in the relationship between changes in fluorescence and behavioral events, we generated the independent variables by convolving a spline basis set with a binary vector of event times (1 at the time of the event, 0 otherwise). The spline basis set was generated with the MATLAB package fDAm.^38^ The full model is:

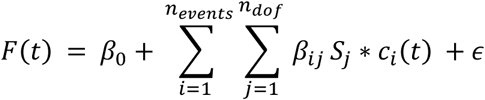

*F*(*t*) is the z-scored ΔF/F at time t, *β*_0_ is the intercept, *ϵ* is the effort, *n_events_* is the number of events, *β_ij_* is the regression coefficient for the *jth S_j_ ith* event, *S_j_* is then *n_dof_jth*, *n_dof_* is the number of degrees of freedom of the basis set and is a binary vector of *i* that is the same length as *F* and is 1 at the time *c_i_i* and 0 otherwise. Convolution of the event vector with a temporal kernel *K_i_*, summed across events:

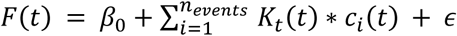

The temporal kernel for event *i* is defined as

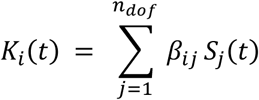

We estimated temporal kernels for the cue and cross events separately for trials when the mouse avoided or escaped the shock, the escape and avoid cross events, and the shock presentation. Temporal kernels for the cue events spanned from 2 seconds before to 1.5 seconds after the event, the shock kernel spanned from 0.5 seconds before to 1.5 seconds after the event, the avoid cross kernel spanned from 1 second before to 5 seconds after the event and the escape cross kernel spanned from 0 seconds to 5 seconds after the event. The degrees of freedom for the basis set was dependent on the duration of the event. Basis sets for cue events had 25 degrees of freedom, shock 14, avoid crosses 42 and escape crosses 35. We estimated these kernels with linear regression with lasso regularization using the MATLAB function lasso. We first selected a regularization parameter, *λ*. Using 5-fold cross validation, we fit the model using a range of *λ* values. We selected the value of*λλ* that minimized the mean squared error calculated with the test data set (∼20% of trials in each cross-validation partition) in each partition and used the average to fit the full data set.

Model performance was assessed by computing the correlation coefficient between the real ΔF/F and the ΔF/F estimated with the model. To estimate confidence intervals, we bootstrapped the dataset by trial 5000 times to obtain a distribution of model fits. To assess the change in goodness of fit across days (Figure 4C), we fit a line to the median of the bootstrapped correlation coefficients by day. To quantify the contribution of the shock event to the model fit (Figure 4D), we fit a second model to the data with the shock predictors removed and then computed the difference in fit (correlation coefficients) for the full model and the reduced model. The removal of the shock event was considered significant if the confidence interval of the difference did not contain 0 (alpha = 0.05 with a Bonferroni correction for multiple comparisons). To compare the amplitude of the response to the cue preceding shock avoidance and escape, we calculated the area under the curve for the estimated kernels excluding the 2 second baseline period using the MATLAB function trapz. We estimated a distribution of AUC from the bootstrapped data and determined that the kernels were significantly different if the confidence interval for the difference in AUC between the events did not contain 0 (alpha = 0.05 with a Bonferonni correction for multiple comparisons).

To quantify changes in event encoding across days (Figure S3), we fit separate models for each mouse and session and calculated the correlation coefficient between the real and estimated ΔF/F for the full model and for reduced models without predictors associated with each event. We then performed a one-way, repeated measures ANOVA on the Fisher z transformation of the correlation coefficients or difference in correlation coefficients.

#### Deep Learning Classification Decoder

To decode neural correlates associated with behavioral outcomes, we developed a deep learning classifier using an architecture composed of Long Short Term Memory and Fully Convolutional Networks (LSTM-FCN) as previously described.^39^ All dopamine recordings for trials occurring on Days 5-7 were used to develop the classifier. Optimal model parameters were determined through a Bayesian hyperparameter optimization.^40^ Hyperparameter sweeps were performed using a dataset split ratio of 0.75 and 0.25 for training and testing respectively. Additionally, all models were trained on the same dataset splits for accurate comparisons. Hyperparameter search space was defined as described (Tables S1, S2) and algorithmically optimized to maximize accuracy on the test dataset. Each model was recorded and analyzed using CometML.^41^ The optimal model was then chosen based on test data accuracy.

#### Other statistical methods

Statistical analysis was done using Prism 10 software (GraphPad). One-way ANOVAs or mixed effects analyses comparing NAc region, day, avoid vs escape, or performance level were performed with Tukey’s multiple comparisons tests when statistically significant main effects or interactions were found (p<0.05). Correlations were conducted using simple linear regressions.

#### Exclusions

One mouse was excluded from the Core experiments due to failure to reach more than 40% avoidance after 7 days of active avoidance testing. Four mice were eliminated due to poor photometry signal, reflected by poor dLight1.3b expression in histology. One mouse was excluded from the Sox 6 experiments due to improper fiber placement and two were excluded due to failure to reach more than 40% avoidance after 7 days of active avoidance testing. All n values listed above do not include these mice.

## Results

### Rapid Acquisition of an Active Avoidance Task with Stable Reaction Times

To determine how NAc dopamine signals contribute to active avoidance learning, we expressed the fluorescent dopamine sensor dLight1.3b in the NAc core (Core) or ventromedial shell (vmShell) and implanted an optical fiber for *in vivo* fiber photometry recordings during freely moving behavior (Figures 1A-B, S1). After recovery from surgery, we trained mice to avoid a footshock punishment (0.4 mA) by moving to the opposite side of a 2-chamber apparatus during a five-second warning cue. Movement to the opposite side of the chamber within 5 s of cue start resulted in cessation of the cue, indicating safety, and avoidance of the shock (“avoid”). If the mouse did not move to the opposite side of the chamber within 5 s, shock punishment began and continued until the mouse moved to the opposite side of the chamber (“escape”; Figure 1C). Mice completed 30 trials per day for 7 days, within random intertrial intervals averaging 45 s. Performance was measured by calculating the percentage of shocks avoided for each day of training. Mice rapidly learned to avoid shocks, reaching an average of 85% avoidance by Day 7 (85.10% ± 7.79; Figures 1D, S2A). The latency to cross to the opposite chamber after the start of the cue also decreased over days (repeated measures one-way ANOVA; F (3.764, 131.7) = 91.49, p< 0.0001), rapidly reaching an average of <5 s, the time required to achieve avoidance (Figures 1E, S2B). This average decrease in latency to cross is closely related to the percent avoidance, but we wondered whether latencies to cross continued to decrease after animals learned the 5s threshold for avoidance. We separated latency-to-cross values by avoid vs escape trials and found that, when the latencies were split by trial type, they remained remarkably stable after Day 1 (Figure 1F). On escape trials, mice quickly reacted to shock initiation and crossed to the opposite chamber within ∼1s (0.86s ± 0.95). On avoid trials, mice consistently crossed at ∼3 s post-cue (2.91s ± 0.32). This stable bifurcation of latencies to cross is due to a discrete shift in latencies over days. While mice crossed predominantly at 5-6 s post-cue during early days of training as a reaction to the shock outcome, they display a bimodal shift to towards 2-3 s latencies over days (Figure 1G). This analysis suggests that mice do not slowly shift their reaction times to earlier points to find the exact latency for shock start, but rather establish a comfortable reaction time that results in successful avoidance and follow a stable avoidance strategy.

**Figure 1.**
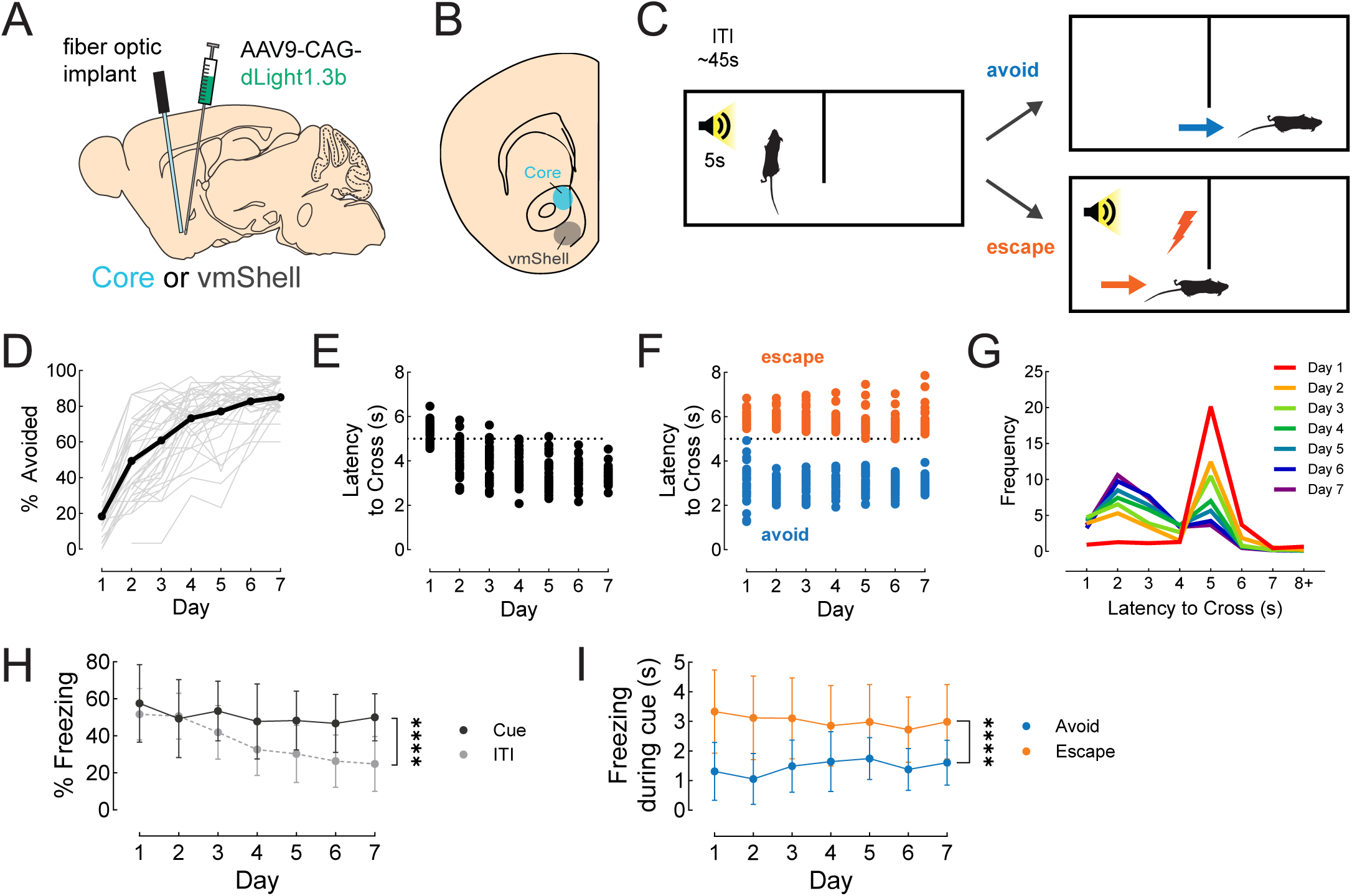
Rapid acquisition of active avoidance with stable reaction times. (A) Mice were injected with an AAV to express dLight1.3b in the NAc Core or vmShell and a fiber optic was placed at the same site to allow collection of fluorescent signals by fiber photometry. (B) Schematic showing the targeted areas for NAc Core (blue) and NAc vmShell (gray). (C) Schematic of the active avoidance task. A 5-s warning cue was played. If the mouse crossed into the opposite chamber before 5s elapsed, the cue turned off and the mouse avoided receiving a shock on that trial (avoid trials). If the mouse did not cross into the opposite chamber before 5s elapsed, shock began. Shock ended when the mouse crossed into the opposite chamber (escape trials). Thirty trials were given per day, with an average inter-trial interval of 45s. (D) Percentage of shocks avoided over days of active avoidance training (n = 36). The black line shows the average for all mice in the study. The gray lines represent the performance of individuals. (E) Latency to cross over days of active avoidance training (n = 36). Each dot is the average for the day from an individual mouse. The dotted line at 5s indicates the latency at which shock would begin. (F) Latency to cross for escape trials (orange) and avoid trials (blue). Each dot is the average for the day from each trial type for an individual mouse (n = 36). The dotted line at 5s indicates the latency at which shock would begin. (G) Frequency plot of average trial latencies for all mice (n = 36). Colors indicate days and show a bimodal shift in distribution from 5-6s to 2-3s over training. (H) Percentage of freezing during the cue (black line) and ITI (grey dashed line) over days of active avoidance (n = 25). Each black square (cue) or gray circle (ITI) represents the average percent freezing of all mice for that day. Percentage of freezing between cue and ITI are significantly different (****p <0.0001). (I) Freezing duration for escape trials (orange) and avoid trials (blue) over days of active avoidance (n = 25). Each dot represents the average freezing duration for all mice for that day. Freezing duration during the cue in avoid vs escape trials are significantly different (****p<0.0001). See also Figures S1 and S2

Since cue-shock pairings can induce freezing as a fear response, we analyzed behavioral videos to quantify freezing during the warning cue and intertrial interval (ITI) periods across training. This analysis showed a main effect of day (mixed effects analysis; F (6, 294) = 17.22, p<0.0001) – meaning mice freeze during the task, but decrease their freezing across days – and a main effect of cue vs ITI period (mixed effects analysis; F (1, 49) = 19.61, p<0.001) with a significant interaction between the two factors (mixed effects analysis, F (6, 147) = 9.845, p<0.0001, Figures 1H, S2C). Freezing significantly decreased across days for the ITI period (correlation; r = -0.97, p = 0.0003), indicating that mice learned the ITI period was safe. Freezing trended lower across days for the cue period (correlation; r = -0.67, p = 0.099), but remained significantly higher than for the ITI period, indicating that mice still displayed fear responses to the cue throughout training. To assess if freezing responses might therefore compete with the performance of avoidance actions during the warning cue, we compared freezing duration during the cue in avoid vs escape trials (Figure 1I) and found a significant difference (mixed effects analysis; F (1, 1115) = 265.6, p<0.0001), indicating that a major reason mice fail to avoid shocks on escape trials is because of their competing intuition to freeze. Taken together, these data show that the mice learn the cue-shock association robustly and then mostly overcome initial freezing responses to learn adaptive active avoidance behaviors by negative reinforcement.

### NAc Core and vmShell have distinct dopamine responses during active avoidance learning

To assess region-specific dopamine signals during active avoidance learning, dopamine release dynamics in the NAc Core and vmShell were recorded across all 7 days of the task. We hypothesized that NAc dopamine signals would show dynamic responses to the warning cue that change with learning. NAc Core and vmShell dopamine signals aligned to the warning cue were distinct and evolved across days (Figures 2A-C, S2D-J). We quantified the area under the curve (AUC) for Core and vmShell dopamine signals during the cue (0-5s) and found a highly significant main effect of brain region (mixed-effects analysis; F (1, 20) = 47.77, p < 0.0001) and day (mixed-effects analysis; F (3.348, 65.85) = 5.860, p = 0.0009), with a significant interaction between the two factors (mixed-effects analysis; F (6, 118) = 9.285, p<0.0001; Figure 2D).

**Figure 2.**
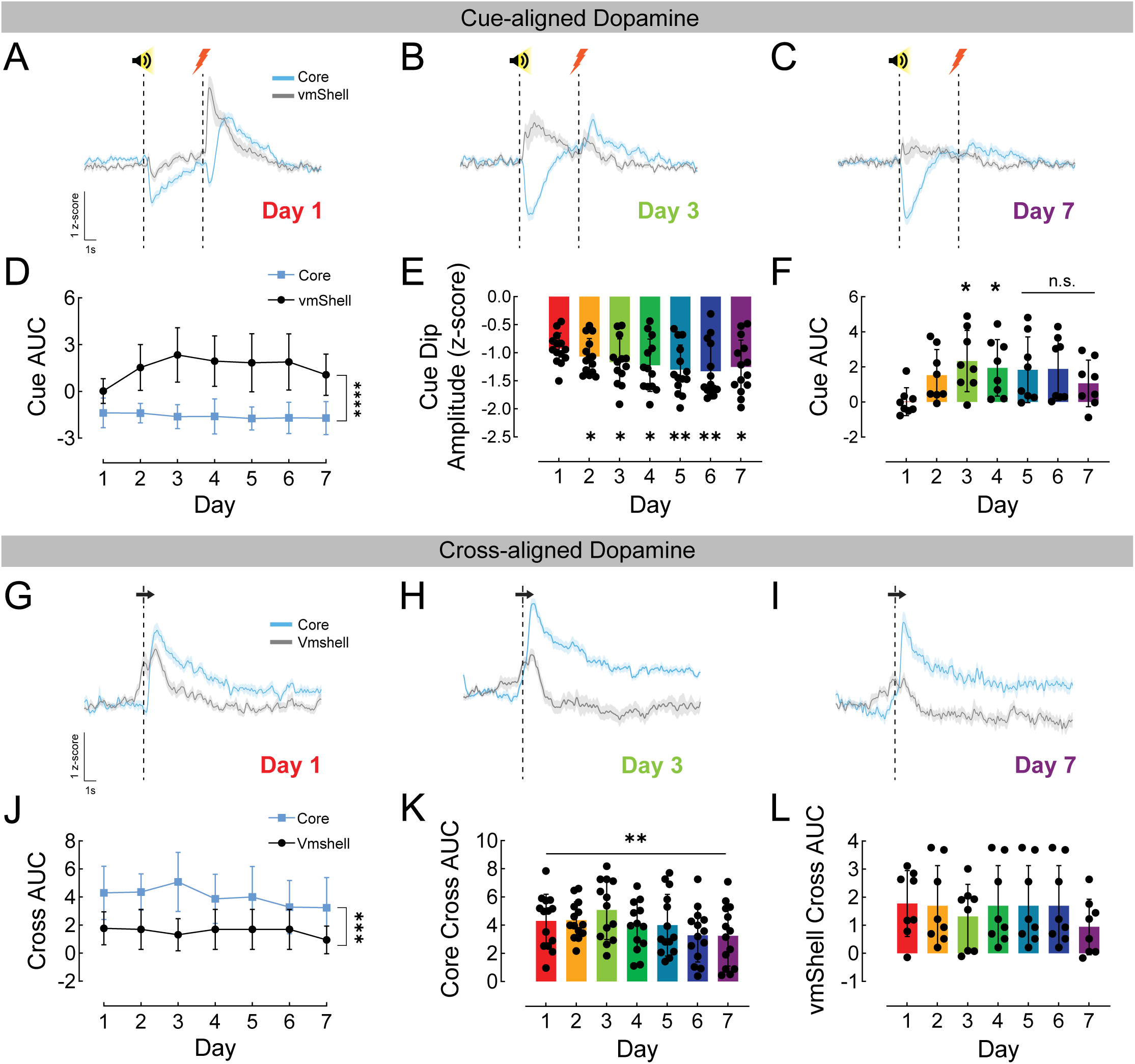
NAc Core and vmShell have distinct dopamine responses during active avoidance learning. (A-C) Plots showing the dopamine signals collected from Core (blue) and vmShell (gray) aligned to the start of the warning cue (first dotted line). Shock occurred 5s after the warning cue start on escape trials (second dotted line). Day 1 (A), Day 3 (B), and Day 7 (C) are shown. (D) The area-under-the-curve (AUC) for the cue period (0-5s) is shown for Core (blue) and vmShell (gray) across days. Cue AUC for Core and vmShell are significantly different (****p<0.0001). (E) Peak negative-going amplitude (cue dip amplitude) for Core over days. Day 2-7 are significantly different from Day 1. *p<0.05, **p<0.01. (F) Cue AUC for vmShell over days. Days 3-4 are significantly different from Day 1. *p<0.05 (G-I) Plots showing the dopamine signals collected from Core (blue) and vmShell (gray) aligned to time of crossing to the opposite chamber (dotted line). Day 1 (G), Day 3 (H), and Day 7 (I) are shown. (J) The AUC for the post-crossing period (0-5s) is shown for Core (blue) and vmShell (gray) across days. Cross AUC for Core and vmShell are significantly different (***p<0.001). (K-L) Cross AUC for Core (K) and vmShell (L) are shown across days. Cross AUC for Core significantly changes across days (**p<0.01). No significant changes are seen across days for vmShell. Core n = 14, vmShell n = 8 See also Figure S2

Core dopamine showed negative-going responses (“dips”) to the warning cue. The amplitudes of these dips grew larger over days (mixed-effects analysis; main effect of day, F (2.304, 29.18) = 11.39, p = 0.0001; Figure 2E). In contrast, vmShell dopamine signals showed an oppositely signed and more complex pattern of change. The cue AUC for vmShell dopamine was minimal on Day 1, grew over the first few days of training, and then decreased later in training. For cue-aligned vmShell dopamine, we found a significant main effect of day, with differences from Day 1 appearing on Days 3 and 4 (repeated measures one-way ANOVA; F (2.550, 17.85) = 7.221, p = 0.0031; Tukey’s multiple comparisons, day 1 versus day 3, p = 0.0208, day 1 versus day 4, p = 0.0452; Figure 2F). These data indicate that Core and vmShell not only have distinct responses during active avoidance, but these responses evolve in unique ways across days of training.

To avoid or escape a shock, mice had to cross to the opposite side of the chamber. Given the importance of this action and dopamine’s role in motivated behavior, we assessed Core and vmShell dopamine release dynamics aligned to crossing. Dopamine signals for both the Core and vmShell increased at the time of crossing (Figure 2G-I). However, Core dopamine was better aligned to the crossing event, whereas as vmShell dopamine began to increase before crossing. Overall, dopamine AUC at the time of crossing (0-5s post-cross) differed by brain region (mixed-effect analysis; F (1, 20) = 16.24, = p 0.0007). We also found a main effect of day (mixed-effects analysis; F (4.423, 86.99) = 2.506, p = 0.0424; Figure 2J). Examining Core and vmShell separately, we found a main effect of day for the Core (mixed-effects analysis; F (4.015, 50.86) = 4.147, p = 0.0055; Figure 2K) but not for the vmShell (repeated measures one-way ANOVA; F (1.533, 10.73) = 0.9050, p = 0.407; Figure 2L), suggesting that Core dopamine specifically is involved learning the value of the avoidance action over days. These differences in dopamine release dynamics for the Core and vmShell propelled our interest in deciphering whether and how these two dopamine signals pertain to different aspects of avoidance learning and performance.

### NAc Core warning cue response dynamics differ by trial type while vmShell dopamine is excited by aversive outcomes

While both avoid and escape events can be characterized as flight behaviors, they differ by a key feature – avoidance eliminates exposure to the shock while escape does not. Some previous work suggested that cue period Core dopamine might promote successful avoidance.^15–16,19^ Given this observation, we sought to decipher how dopamine signals in the Core and vmShell differ between avoid and escape trials in our task (Figure 3). When aligned to the cue, Core dopamine release dynamics differed by trial type (avoid vs escape; mixed-effects analysis; F (1.000, 13.00) = 94.17, p<0.0001, Figure 3A-D). There was also a main effect of day (mixed-effects analysis; F (3.800, 49.40) = 3.767, p = 0.011) indicating changing signals across learning. In contrast, vmShell dopamine dynamics aligned to the cue did not differ by trial type (mixed-effects analysis; F (1.000, 7.000) = 1.845, p = 0.2165, Figure 3E-H). We did, however, observe vmShell dopamine responses on escape trials in response to shock onset, indicating that vmShell dopamine is excited by aversive stimuli (Figure 3E-G).

**Figure 3.**
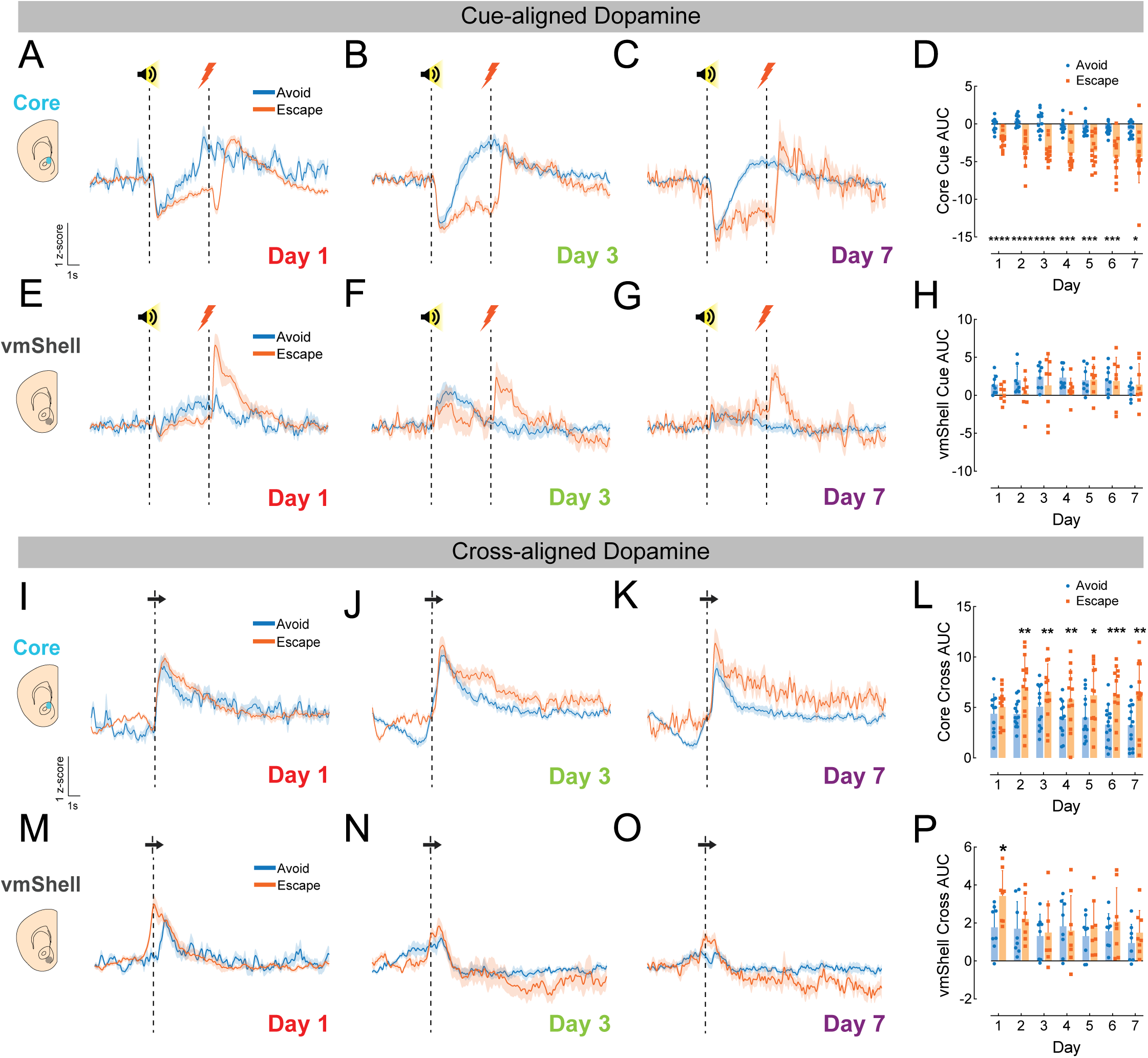
NAc Core and vmShell dopamine responses differ by trial type. (A-C) Plots showing Core dopamine signals collected during avoid trials (blue) and escape trials (orange) from aligned to the start of the warning cue (first dotted line). Shock occurred 5s after the warning cue start on escape trials (second dotted line). Day 1 (A), Day 3 (B), and Day 7 (C) are shown. (D) The Core area-under-the-curve (AUC) for the cue period (0-5s) is shown for avoid and escape trials across days. Cue AUC for avoid vs escape trials are significantly different across days *p<0.05, ***p<0.001, ****p<0.0001. (E-G) Plots showing vmShell dopamine signals collected during avoid trials (blue) and escape trials (orange) from aligned to the start of the warning cue. Day 1 (E), Day 3 (F), and Day 7 (G) are shown. (H) The vmShell AUC for the cue period (0-5s) is shown for avoid and escape trials across days. No significant differences are observed between avoid and escape trials. (I-K) Plots showing Core dopamine signals collected during avoid trials (blue) and escape trials (orange) aligned to time of crossing to the opposite chamber (dotted line). Day 1 (I), Day 3 (J), and Day 7 (K) are shown. (L) The Core AUC for the post-crossing period (0-5s) is shown for is shown for avoid and escape trials across days. Avoid and escape cross AUCs are significantly different from each other on Days 2, 5, 6, and 7 (*p<0.05, **p<0.01, ***p<0.001). (M-O) Plots showing vmShell dopamine signals collected during avoid trials (blue) and escape trials (orange) aligned to time of crossing to the opposite chamber (dotted line). Day 1 (M), Day 3 (N), and Day 7 (O) are shown. (P) The vmShell AUC for the post-crossing period (0-5s) is shown for is shown for avoid and escape trials across days. A significant difference between avoid and escape trials is observed on Day 1 (*p<0.05). Core n = 14, vmShell n = 8 See also Figure S4

We also examined how dopamine signals for avoid and escape trials differed when aligned to crossing. In the Core, cross-aligned dopamine dynamics during avoid vs escape trials slowly diverge over days (Figure 3I-K). We found a significant main effect of trial type (mixed-effects analysis; F (1.000, 13.00) = 35.39, p<0.0001) with significant differences between avoid and escape trials emerging after Day 1 (Figure 3L). When vmShell cross AUC was quantified across days, we found no significant effect of trial type, but a significant main effect of day (repeated measures two-way ANOVA; F (4.010, 28.07) = 3.234, p = 0.0265; Figure 3M-P).

To better understand how behavioral events predicted features of the fiber photometry data, we built linear encoding models^42–45^ for each brain region and day of training to estimate temporal kernels modeling responses to cues, shocks, and crossings (Figure 4A-B). This analysis provoked several additional insights. We found that the goodness of fit for the overall model for vmShell dopamine worsened over days (slope: -0.0327, p = 0.008), indicating that vmShell dopamine representations of task events weaken with time (Figure 4C). To measure the contribution of individual events to dopamine activity, we compared the fit of the full model to a reduced model with each event type removed. Using this approach to evaluate shock encoding, we found that the shock event contributed significantly to vmShell dopamine across all days, but that the relationship was strongest on Day 1, and weakened by Day 3 (Figure 4D). We verified that shock encoding in the vmShell dopamine signal changed across days by modeling dopamine fluorescence in individual mice and assessing the influence of the shock event on model performance. We found that shock representation decreased across days (one-way, repeated measures ANOVA, F (6,36) = 4.88 p = 9.64x10^-4^; Figure S3). This decrease was not only due to the worsening of the full model for vmShell over time, as representations of other events did not decrease over days (Figure S3). These observations suggest that vmShell might be important for early learning about shock consequences, but less so for consolidating and sustaining avoidance. In contrast, the overall model for Core dopamine remained strong across days (slope: -0.004, p = 0.437; Figure 4C), with robust encoding of warning cues and avoidance actions (Figure 4A). Interestingly, the Core dopamine model showed strengthening representations of warning cues on avoid trials compared to escape trials as learning progressed, with the kernels for avoid vs escape trials diverging on Days 5-7 (Figure 4E).

**Figure 4.**
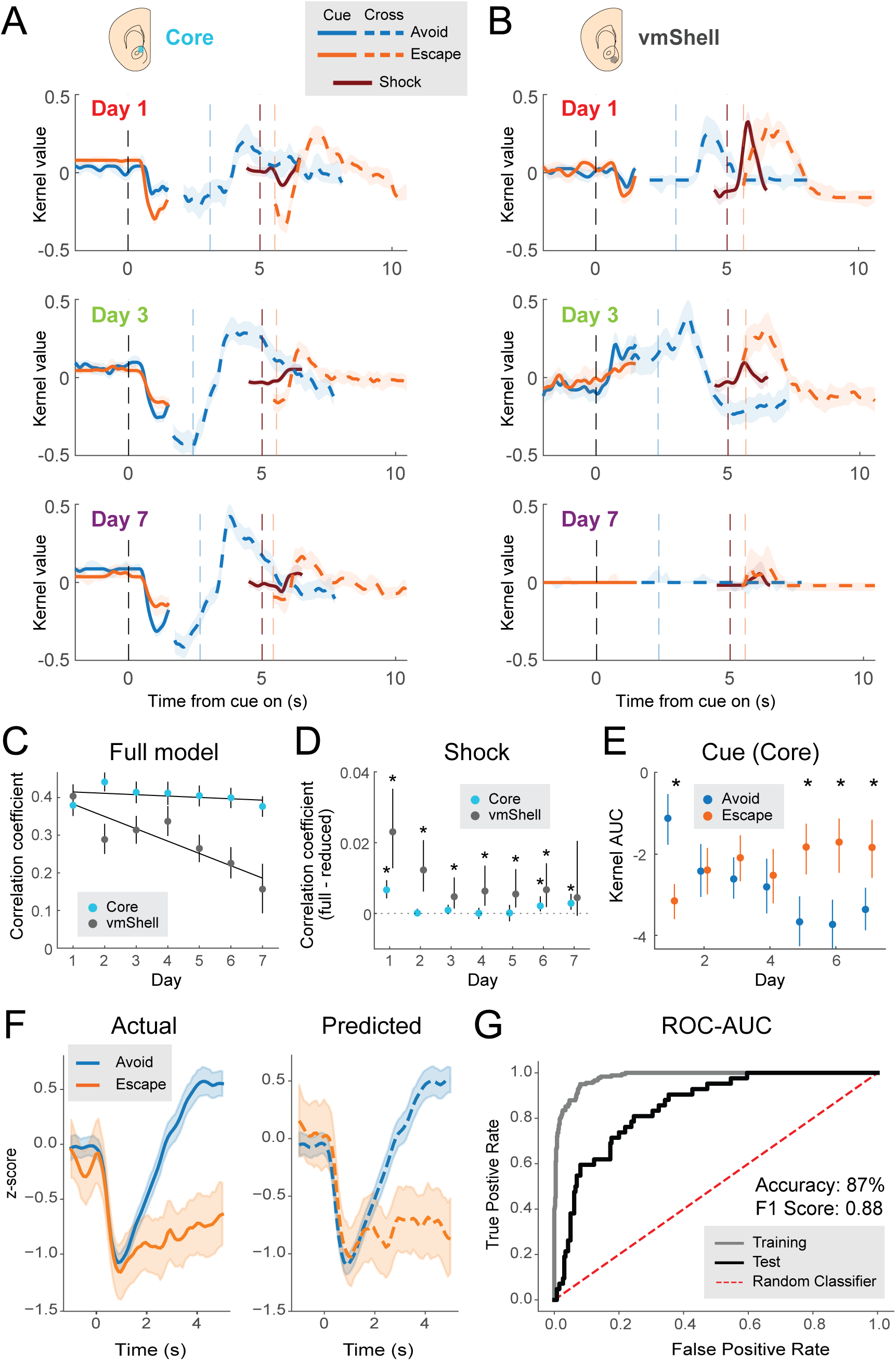
Distinct dynamics of NAc Core and vmShell dopamine encoding over learning. (A-B) Kernel values from the encoding model of the photometry signals for Core (A) and vmShell (B) are shown for Days 1, 3, and 7 of training. Behavioral events are marked by vertical dashed lines: cue-on (black), median crossing time for avoid trials (blue), median crossing time for escape trials (orange), shock-on (for escape trials only; dark red). (C-E) Data points are the median of the bootstrapped distribution. Error bars are 95% confidence intervals. (C) Correlation coefficients over days for the full encoding models for Core and vmShell dopamine. The rate of change is estimated by fitting a line to each set of data points (Core slope = -0.004, p = 0.437; vmShell slope = -0.0327, p = 0.008). (D) The change in correlation coefficients when shock is removed as a behavioral event (full model – reduced model). Asterisks indicate that the difference in correlation coefficients is significantly different from 0 (based on confidence intervals with a Bonferroni correction for multiple comparisons). (E) Area under the curve (AUC) for the kernel values for avoid-cue and escape-cue over days. Asterisks indicate that the difference in AUC for the two events is >0 (based on confidence intervals with a Bonferroni correction for multiple comparisons). (F) Core dopamine signals during the cue period for avoid (blue) and escape (orange) trials on Days 5-7. True signals are shown on the left (solid lines) and signals predicted to be avoid vs escape using the LSTM-FCN classification decoder are shown on the right (dashed lines). (G) Receiver operator characteristic (ROC) curve for the LSTM-FCN classification decoder. The gray line represents the ROC of the training dataset, the black line represents the ROC of the test dataset, and the red dashed line represents random classification. See also Figure S3 and Tables S1-3

This finding prompted us to test if decoding avoid and escape trials from Core dopamine signals during the cue period on Days 5-7 might also be possible. We implemented a Long Short-Term Memory Fully Convolutional Network (LSTM-FCN),^39^ which allowed us to model noisy signals without additional feature extraction or manipulation. We optimized model parameters using a Bayes optimization algorithm to maximize prediction accuracy. On test data, our decoding model achieved 87% accuracy (Figure 4F-G, Table S3). Furthermore, accounting for label imbalances, we achieved an F1 score of 0.88. Lastly, analysis of the receiver operator characteristic (ROC), a common metric examining the relationship between true and false-positive rates, demonstrated an area under curve (ROC-AUC) of 0.85. These results verify that the information contained within the Core dopamine signal is largely sufficient to decode established avoidance behavior on a trial-by-trial basis following learning.

### NAc vmShell dopamine responses to the warning cue evolve rapidly during early avoidance learning

Prompted by our modeling results showing that Core and vmShell dopamine evolve differently across learning, we sought to better understand the dynamics of each signal separately. VmShell dopamine representations of the shock were high early and faded quickly. VmShell dopamine representations of the warning cue rose during early days of avoidance learning (as shock responses faded), but also were not sustained, fading on Days 4-7 (Figures 2F, 1D). We hypothesized that vmShell dopamine cue responses are linked to acquisition of the cue-shock association but are not required for avoidance performance. To address our hypothesis, we began by analyzing learning rates more closely, binning avoidance performance in 10-trial blocks over days (Figure 5A). Examining the percentage of shocks avoided across 10-trial bins revealed that the largest learning gains occurred during Days 1-3, with relatively stable performance across the session on Days 4-7. We next examined within-day changes on Days 1-3 even more closely, plotting the percentage of mice avoiding the shock on each of the 30 trials for these days (Figure 5B). We observed a slow, gradual increase in performance on Day 1. Although some “forgetting” occurred overnight, leading to poorer performance on the first few trials of Days 2 and 3, we observed rapid performance improvements over trials on these days. Consistent with our hypothesis, vmShell dopamine signals changed dramatically on Days 1-3, shifting from the shock to the cue (Figure 5C). To explore within-day changes during learning, we quantified the vmShell cue AUC for Days 1-3 across 10-trial bins. However, we found no main effect of trial bins within days (two-way ANOVA; F (1.897, 13.28) = 0.6054, p = 0.552, Figure 5D), indicating this shift is not tied closely to performance metrics within days.

**Figure 5.**
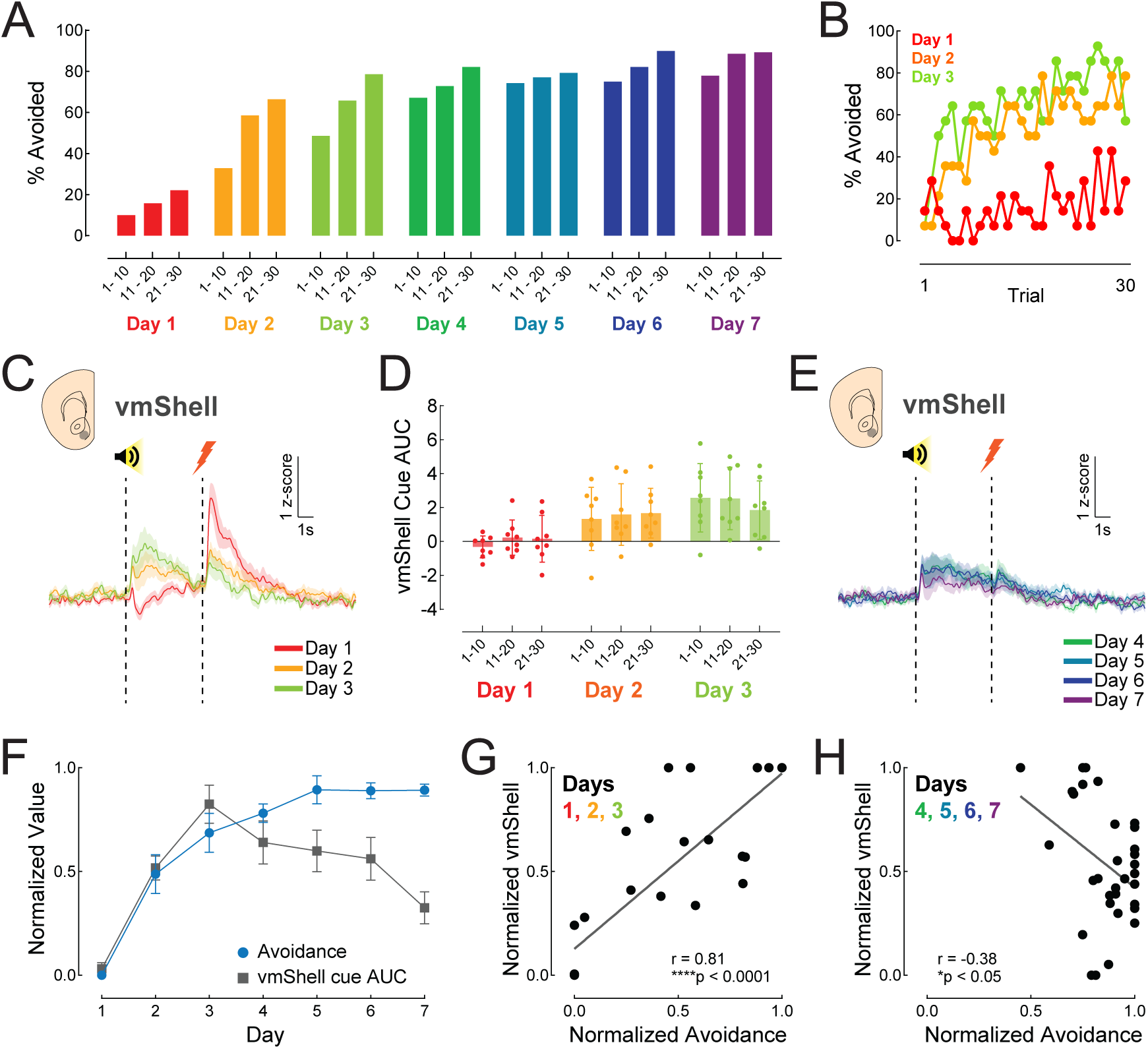
NAc vmShell dopamine responses to the warning cue evolve rapidly during early avoidance learning. (A) Percentage of shocks avoided during 10-trial blocks over days of active avoidance training. Large learning gains occurred on Days 1, 2, and 3. Asymptotic performance was observed on Days 4, 5, 6, and 7. (B) Percentage of shock avoided by trial number on Days 1 (red), 2 (orange), and 3 (green). (C) Cue-aligned dopamine signals recorded from vmShell on Days 1 (red), 2 (orange), and 3 (green) change dramatically across learning days. (D) The vmShell AUC for the cue period (0-5s) is shown for 10-trial blocks across learning days (1, 2, and 3). (E) Cue-aligned dopamine signals recorded from vmShell on Days 4 (dark green), 5 (teal), 6 (navy) and 7 (purple) remain steady during asymptotic performance. (F) Percentage of shocks avoided (blue) and vmShell cue AUC (gray) are shown as normalized values for comparison (0=lowest value in dataset; 1=highest value in dataset) across days of avoidance learning. (G) Cue-period vmShell dopamine scales with the acquisition of avoidance behavior on learning days (r = 0.81, ****p<0.0001) (H) Cue-period vmShell dopamine is negatively correlated with the performance of avoidance behavior after learning r = 0.38, *p<0.05 VmShell n = 8

In contrast to the dramatic changes in cue-aligned vmShell dopamine during Days 1-3, the signals on Days 4-7 decreased gradually in response to the cue (Figure 5E). To compare changes in learning and changes in vmShell dopamine signaling across days, we normalized both of these measures for each mouse, with 0 being the lowest within-subject value and 1 being the highest (Figure 5F). This visualization makes clear that vmShell cue dopamine rises in step with early task acquisition but falls off from performance later. Quantitatively, we found a strong, highly significant positive correlation between normalized avoidance and normalized vmShell cue dopamine for Days 1-3 (r = 0.81, p < 0.0001; Figure 5G), and a weak negative correlation for days 4-7 (r = -0.38, p<0.05; Figure 5H). These data are all consistent with the idea that although cue period dopamine in the vmShell is altered during early avoidance learning, it is unlikely to contribute to performance consolidation as subjects solidify their understanding of the context-specific avoidance rules.

### NAc Core dopamine responses to the warning cue reflect consolidation of avoidance learning at high performance levels

Changes in NAc Core dopamine responses to the warning cue follow a very different temporal pattern than vmShell dopamine responses. In Core, there is a dip in dopamine that deepens slowly across days of training and is well-linked to successful avoidance in late training (Figures 3-4). We hypothesized that this deepening dip was related to the consolidation of avoidance performance. To assess this hypothesis, and to differentiate performance from time in training, we re-analyzed our Core dopamine data by individual performance level rather than by day.

Even though most mice rapidly learn to avoid shocks across days of training, there is variability in the rate at which an individual mouse reaches asymptotic, expert performance (Figure 1D, gray lines represent individuals). Given this variability, we decided to look at the distribution of mice achieving defined levels of performance across days of training. We analyzed when mice reached threshold of >25%, >50%, and >75% performance (Figure 6A). The resulting histogram shows a wide distribution of days when expert performance (>75% avoidance) is achieved, with some mice reaching this expert level only on the final day of training and one mouse never reaching this level. Comparing this performance data to Core dopamine signals, we found that Core dopamine dip amplitude correlates significantly with performance, with larger dips in the Core cue-aligned dopamine signal predicting higher rates of avoidance (correlation; r = -0.46, p < 0.0001; Figure 6B). To visualize this relationship, we parsed out Core dopamine signals aligned to the cue by performance levels (0-25%, 26-50%, 51-75%, 76-100%) rather than day of training (Figure 6C). There was a main effect of performance level by this analysis (mixed-effects analysis; F (2.703, 82.88) = 8.210, p = 0.0001). We also observed that the dip gets much larger specifically at the highest performance level (Dunnett’s multiple comparisons; 0-25% versus 76-100%, p = 0.0004; Figure 6D). Thus, in contrast to vmShell cue dopamine signals, dopamine signals in the Core are strongly linked to consolidation of expert avoidance.

**Figure 6.**
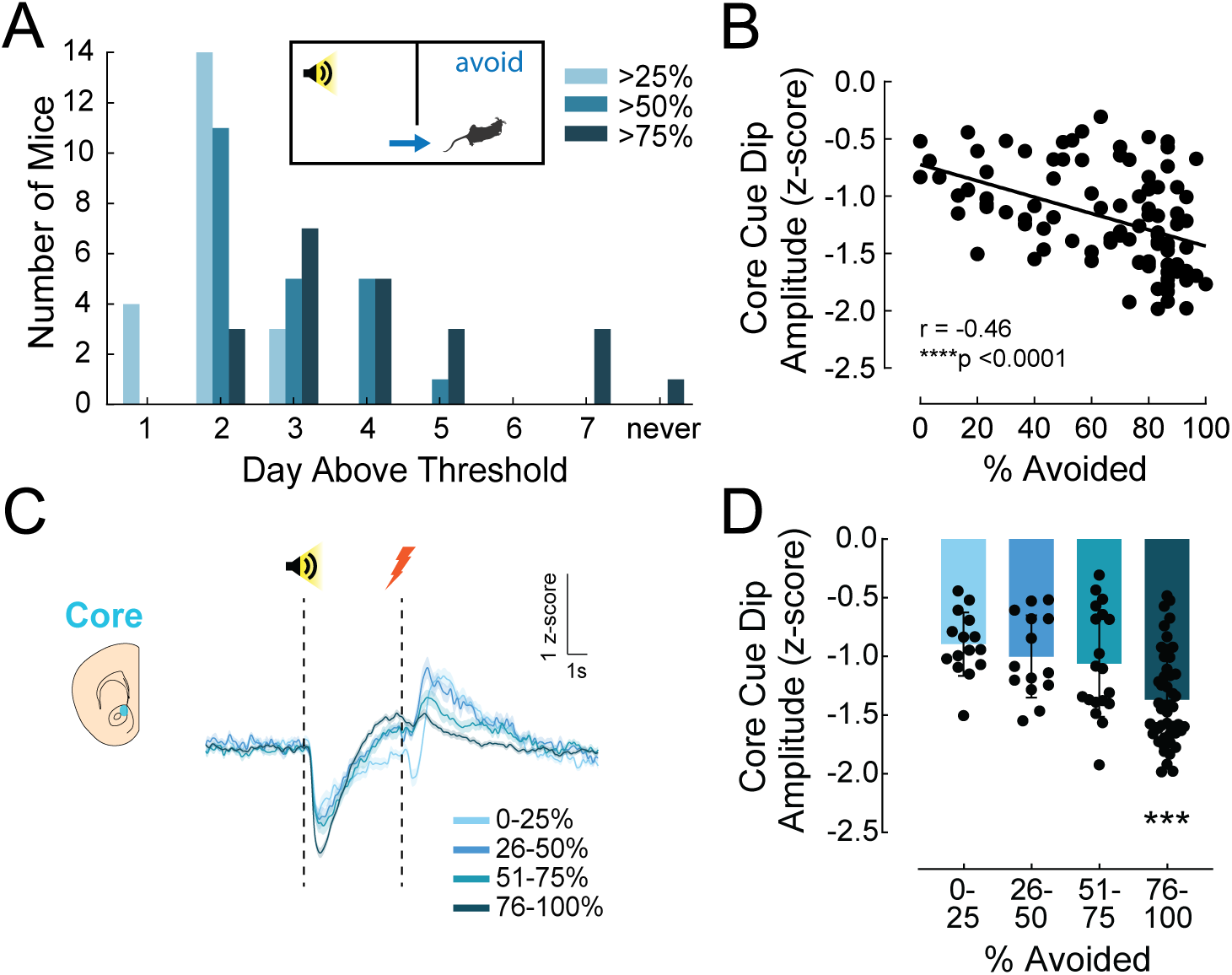
NAc Core dopamine responses to the warning cue reflect the consolidation of avoidance learning at high-performance levels. (A) Histogram showing the number of mice reaching a threshold of performance criteria on each day. Light blue shows the number of mice reaching 25% avoidance or better for the first time on that day, medium blue shows the number of mice reaching 50% avoidance or better for the first time on that day, dark blue shows the number of mice reaching 75% avoidance or better for the first time on that day. Although mice generally learn the avoidance task rapidly, there is individual variability in the speed of learning. (B) Core cue dip amplitude correlates with the percentage of avoided shocks (r = -0.46, ****p<0.0001). (C) Cue-aligned dopamine signals recorded from Core divided out according to the avoidance performance of the mouse rather than day recorded. (D) Peak negative-going amplitude (cue dip amplitude) for Core by performance level. The cue dip amplitude changes minimally at lower performance levels (up to 75% avoidance) but becomes significantly larger on days with high performance (76-100% avoidance; ***p<0.001 compared to 0-25% avoidance). Core n = 14 See also Figure S4

To evaluate how Core dopamine release dynamics during avoidance consolidation influence downstream circuit function in the basal ganglia, we also recorded the activity of dopamine-receptor 1 and dopamine-receptor 2 containing spiny projection neurons (D1-SPN and D2-SPN, respectively) in the Core during active avoidance learning (Figure S4). We expressed the calcium sensor jRCaMP1b in the NAc Core of D1-Cre or A2A-Cre mice and implanted an optical fiber for in vivo fiber photometry recordings. For both D1-and D2-SPNs, we observed increases in calcium activity in response to the cue and shock (Figure S4B-D; S4I-K), but there were no differences in cue AUC based on trial type (Figure S4E, L). Given that we saw Core dopamine dynamics evolve across performance, we also plotted D1-and D2-SPN calcium activity in the same fashion (Figure S4F, M). Quantification of cue AUC across performance aligned to the cue and shock showed that D1-SPNs have significant increases early in training for the cue (Cue AUC: Brown-Forsythe ANOVA test; F = 4.114 (3.000, 28.20), p = 0.0154; Tukey’s multiple comparison’s; 26-50% vs 51-75%, p = 0.0243, Figure S4G-H), while D2-SPNs have significant increases late in training for cue and shock (Cue AUC: Brown-Forsythe ANOVA test; F = 3.938 (3.000, 18.47), p = 0.0248; Shock AUC: Welch’s ANOVA test, F = 4.389 (3.000, 18.70), p = 0.0168; Dunnett’s T3 multiple comparison’s; 26-50% vs 76-100%, p = 0.0151, Figure S4N-O). Together these data suggest an interesting recruitment mechanism of downstream targets in Core responses to avoidance learning.

### NAc dopamine responses to escapable shock are distinct from responses to inescapable shock

Our data agree with previous data reporting that Core dopamine decreases and vmShell dopamine increases for shocks and shock-predicting cues.^20^ However, our study adds important information regarding how these opposite dopamine signals are used and change during instrumental learning. During an active avoidance escape trial, even though a mouse endures shock, it can remove the negative stimulus by crossing to the opposite chamber. Therefore, instrumental access to safety is retained (Figure 7A). How does the controllability of the negative stimulus change dopaminergic encoding? To investigate possible differences in dopamine release dynamics between escapable and inescapable scenarios, we exposed mice to inescapable shocks at the end of avoidance training (Day 8). During the inescapable scenario, the same 5s warning cue that was used during avoidance learning was followed by a 5s footshock punishment regardless of the animal’s action (Figure 7B). We repeated the inescapable shock for 10 trials while recording dopamine signals from Core and vmShell.

**Figure 7.**
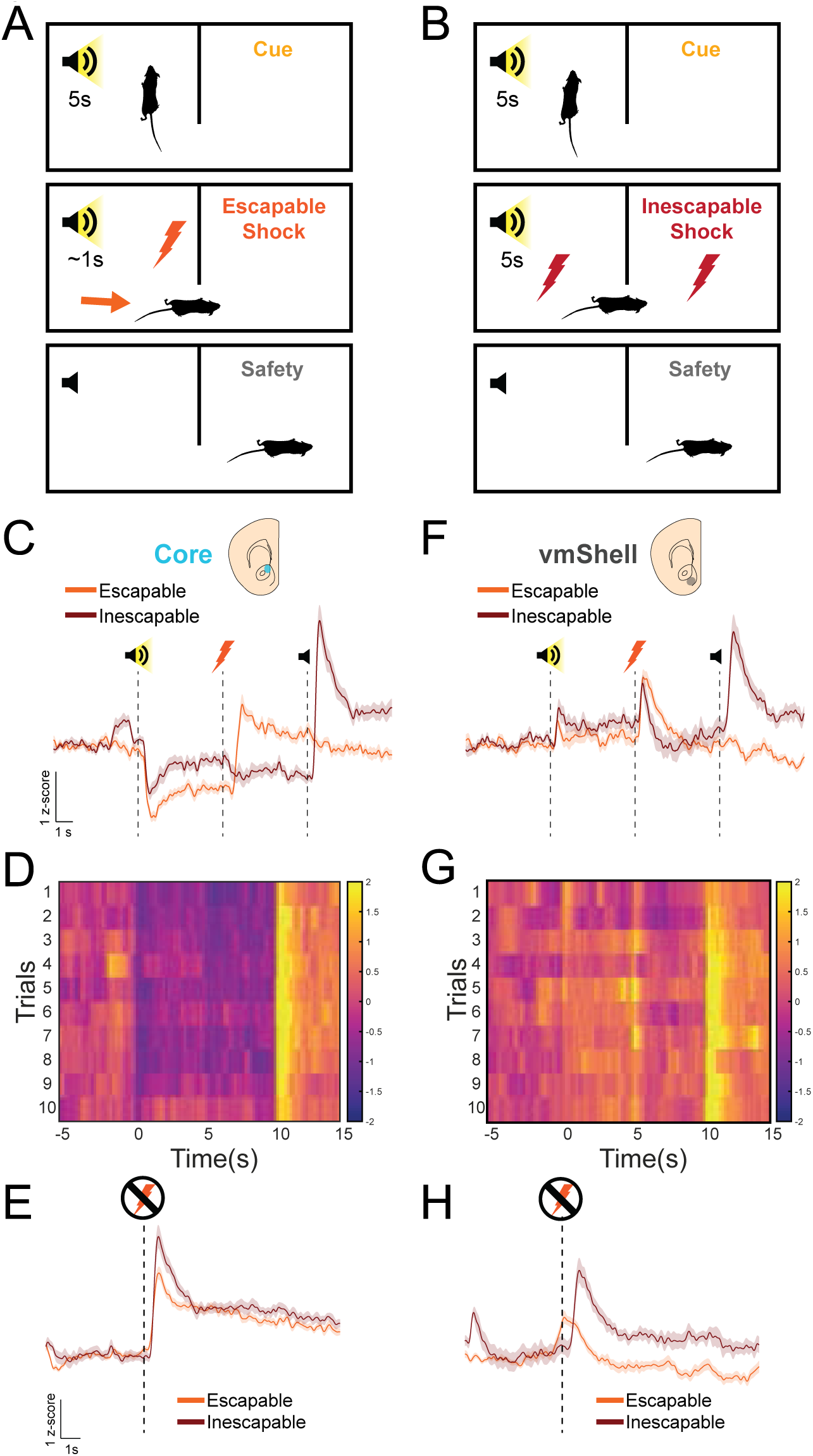
NAc dopamine responses to escapable shock are distinct from responses to inescapable shock. (A-B) Schematics comparing escapable and inescapable shock trials. Escapable shock trials from high performance days (76-100% avoidance days, when mice have learned the avoidance task well) are shown in the data below. On these trials, mice hear a 5s warning cue, shock begins, and mice can escape to safety in the opposite chamber. Escapable trials are compared to a final day of behavioral testing with 10 inescapable shock trials. In these trials, mice hear a 5s warning cue, then experience 5s of shock that cannot be escaped. After 5s, the cue and shock end and the mouse is safe for the inter-trial interval period, which averages ∼45 seconds. (C,F) Cue-aligned dopamine signals recorded in the Core (C) and vmShell (F) for escapable (orange) and inescapable (dark red) shock trials. The first dotted line indicates cue start, the second dotted line indicates shock start, the third dotted line indicates shock end for inescapable shock. (D,G) Heat maps showing the cue-aligned dopamine signals for Core (D) and vmShell (G) on each of the 10 inescapable shock trials (1-10, shown descending in time). (E,H) Dopamine signal for inescapable shock (orange) or escapable shock (dark red) for Core (e) and vmShell (h) in response to controllable or uncontrollable shock termination. The dotted line indicates the time of shock end. Core n = 14, vmShell n = 8

To compare escapable and inescapable shocks, we examined dopamine signals from escape trials of expert performance mice (i.e., trials when mice experienced shocks but were aware of how to escape) and compared these recordings to signals recorded during 10 trials of inescapable shock. When comparing the initial dips in Core dopamine in response to the warning cue, we observed weakened cue-related dynamics in response to inescapable shock (Figure 7C). During inescapable shock trials, Core signal remained below baseline for the entire cue period and further decreased during the subsequent shock. At shock end, the inescapable Core signal showed a large phasic increase. Trial-by-trial analysis of the inescapable Core signal shows relatively similar signal patterns across trials (Figure 7D). To put the large phasic increase at shock-end in context, we compared the signal for the end of inescapable shock to the signal for the end of escapable shock (Figure 7E). The shock-end signal is of larger magnitude when the shock was inescapable, consistent with a greater prediction error.

When comparing escapable to inescapable shock signals in the vmShell, both trial types resulted in phasic dopamine increases in response to cue and shock (Figure 7F). Cue-related dopamine dynamics in the vmShell, which had faded with learning, are strengthened during inescapable shock. Like the inescapable Core signal, the inescapable vmShell signal shows a large phasic increase at shock-end that is consistent or even grows slightly over trials (Figure 7F-G). Comparing the signal for the end of inescapable shock to the signal for the end of escapable shock, we observed very different signaling patterns in vmShell (Figure 7H). Aligned to the end of an escapable shock, vmShell dopamine showed an increase that began a second before the time of shock removal. Given our earlier analysis of this signal (Figure 4) and knowing that mice escape shocks within 1s or less during the avoidance task, we conclude that this signal is a weak shock-start signal. Meanwhile, the vmShell dopamine signal for the end of inescapable shock was time-locked to the shock removal event. This signal is a new dopamine signal that was not observed during avoidance learning (Figure 7H). Previously, we had observed vmShell dopamine responses for aversive events (shocks and shock-predicting cue), but the inescapable shock-end signal might feasibly be related to salience.^12^ These data suggest that dopamine signals can help identify changes in avoidance learning rules, which is important for distinguishing behaviors that may require divergent response strategies for survival. Consistent with our observation that vmShell dopamine is most relevant during early learning, the switch to a new rule during inescapable shock trials increased cue-aligned vmShell dopamine, which had faded during late avoidance training.

### NAc Core dopamine axon activity does not depend on molecular identity and diverges from dopamine dynamics at high performance levels

We observed heterogeneity of dopamine responses during avoidance learning at a projection area level, but we also wanted to investigate whether this regional specificity is based on molecular identity. Previous work focusing on locomotor encoding in substantia nigra pars compacta (SNc) dopamine subpopulations has suggested that molecular differences between subtypes of dopamine neurons can contribute to projection-dependent specificity in dopamine signaling.^46^ The NAc Core is an interesting transition zone between the ventral and dorsal striatum. Ventrally and dorsally projection dopamine neurons can be broadly differentiated by their expression of the transcription factor Sox6.^34^ Sox6+ dopamine neurons populate the ventral tier of SNc and dorsolateral VTA, while Sox6-dopamine neurons populate the dorsal tier of SNc as well as VTA. We observed that the Core, particularly the dorsomedial part of Core where we recorded dLight1.3b signals, receives inputs both from dopamine neurons that express and do not express Sox6 (Figure 8A). It is not clear, however, if these distinct molecular subtypes contribute differentially to Core dopamine signals for avoidance learning. To examine the impact of molecular identity on Core dopamine signals in an active avoidance task, we crossed Dat-2A-Flpo or Th-2A-Flpo mice to Sox6-FSF-Cre,^33,34^ resulting in mice that express Flpo in all dopamine neurons and Flpo + Cre only in Sox6+ dopamine neurons. We injected DAT/TH-2A-Flpo x Sox6-FSF-Cre mice with a virus expressing cre-on/flpo-on-GCaMP6f (AAV-Ef1α-Con/Fon-GcaMP6f) or cre-off/flp-on-GCaMP6f (AV-Ef1a-Coff/Fon-GcaMP6f) to target Sox6+ or Sox6-expressing dopamine neurons, respectively (Figure 8B). A fiber optic implant was placed above the NAc Core to measure VTA dopamine axon activity as mice learned the active avoidance learning task (Figures 8C, S5).

**Figure 8.**
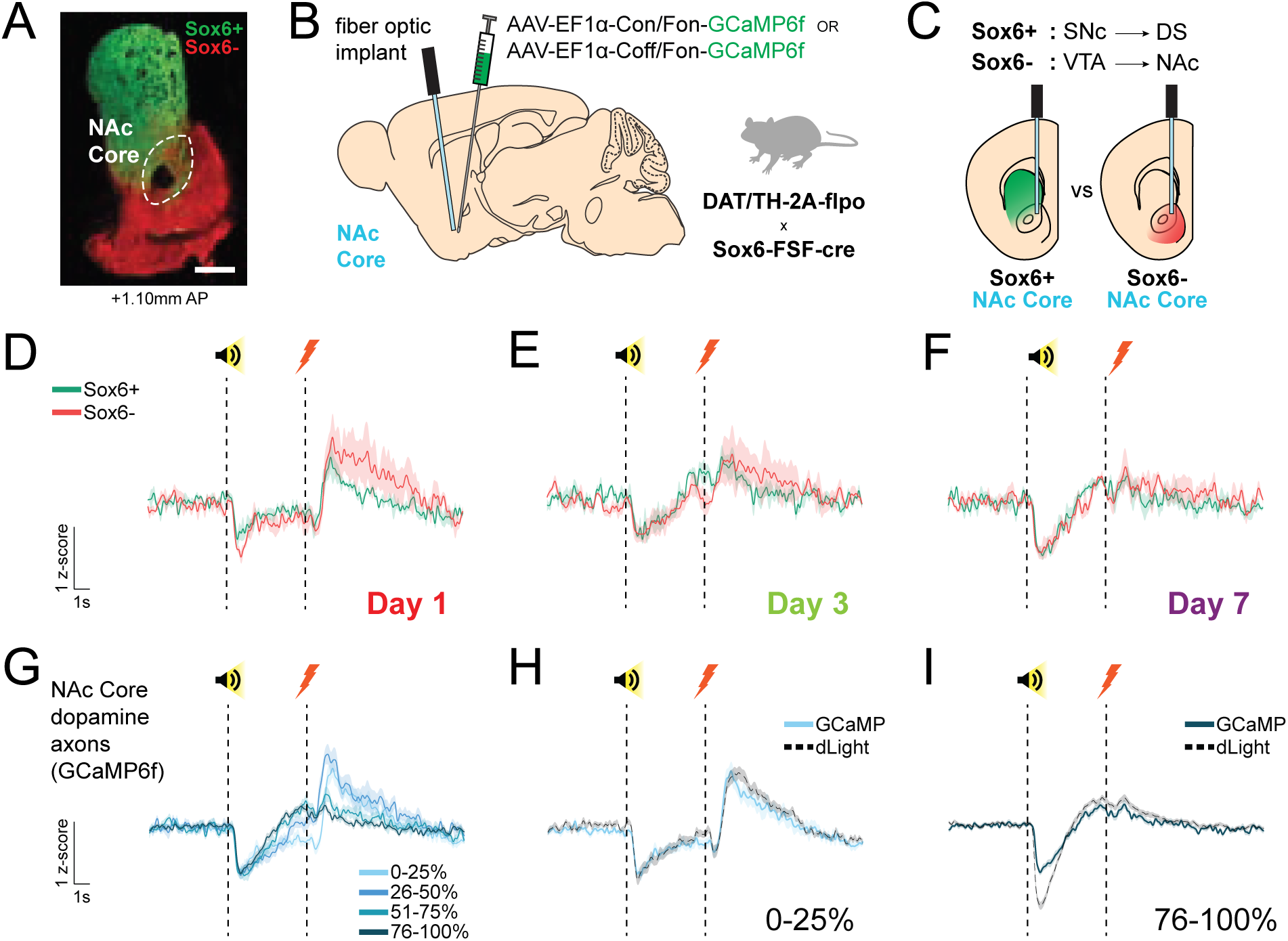
NAc Core dopamine axon activity is independent of molecular identity and diverges from dopamine release dynamics at high-performance levels. (A) Representative histology image showing labeling of Sox6+ and Sox6-dopamine axons in Sox6-FSF-Cre/Dat-2A-Flpo mice. Con/Fon-YFP labeled Sox6+ dopamine axons (green) and Coff/Fon-mCherry labeled Sox6-dopamine axons (red). White dashed line outlines the NAc Core. Scale bar is 250 µm. (B) Schematic showing the experimental strategy for expressing GCaMP6f in Sox6+ or Sox6-dopamine axons. DAT-2A-flpo mice express flpo recombinase in dopamine neurons. This mouse line was crossed to a Sox6-FSF-cre line that expresses cre recombinase in a flpo-dependent manner. Therefore, Sox6+ dopamine neurons express flpo and cre, while Sox6-dopamine neurons express only flpo. Viruses to express GCaMP6f in these combinatorically defined cell types were injected into the NAc Core, and fiber optic implants were placed for fiber photometry. (C) Sox6+ dopamine neurons are generally SNc dopamine neurons projecting to dorsal striatum (DS) while Sox6-dopamine neurons are generally VTA dopamine neurons projecting to NAc. The schematic illustrates these projection zones and shows that fiber photometry probes placed in NAc Core are within a transition zone between the Sox6+ and Sox6-dopamine projection areas. (D-F) Plots showing the GCaMP signals collected from Sox6+ (green) and Sox6-(red) dopamine axons aligned to the start of the warning cue (first dotted line). Shock occurred 5s after the warning cue start on escape trials (second dotted line). Day 1 (D), Day 3 (E), and Day 7 (F) are shown. (G) Cue-aligned GCaMP signals recorded from Core dopamine axons (Sox6+ and Sox6-, pooled) divided out according to the avoidance performance of the mouse. (H) Comparison of Core GCaMP dopamine axon signals and dLight dopamine signals during 0-25% performance. (I) Comparison of Core GCaMP dopamine axon signals and dLight dopamine signals during 76-100% performance. Sox6+ n = 4, Sox6-n = 3 See also Figure S5

Dopamine axon activity in the Core was recorded across the 7 days of active avoidance training. Dopamine axon activity from both Sox6+ and Sox6-dopamine neurons resembled our previous dLight recordings from Core. Recordings from Sox6+ axons did not differ from recordings from Sox6-axons across all days of training (Figure 8D-F). This finding was surprising, but indicates that signals for avoidance learning may depend more strongly on projection region than on molecular identity, or alternatively, other molecular markers may provide more functionally relevant distinctions.

There is controversy on whether measurements of dopamine axon activity via calcium sensors mirror dopamine release dynamics using dopamine binding sensors such as dLight.^47,48^ Despite the lack of differences between Sox6+ and Sox6-dopamine axons in our experiments, we realized that our data could speak to this notion. Given that Core dopamine signals measured with dLight1.3b evolved across performance levels of the avoidance task (Figure 6), we examined pooled (Sox6+ and Sox6-) dopamine axon activity for different performance levels as well. Core dopamine axon activity at outcome evolved across performance levels, but cue-aligned dip amplitudes in dopamine axon activity did not change (Figure 8G). Specifically comparing recordings from mice at low (0-25% avoidance) and high (76-100% avoidance) performance levels revealed that Core dopamine dynamics diverge from dopamine axon activity during high performance (Figure 8H-I). While Core dopamine axon activity overlapped with dLight signals during low performance (Fig. 8H), at high performance (Figure 8I), the cue-aligned dip was greater in magnitude for the dLight signal compared to the GCaMP axon signal. These data indicate that Core extracellular dopamine dynamics may be influenced by factors other than dopamine axon calcium dynamics and that these other factors may become particularly important during learning.

## Discussion

Here, we explored how dopamine signals – well known to support positive reinforcement learning – participate in learning from negative reinforcement. We show that dopamine signals in the NAc Core and vmShell are not only heterogeneous in their responses to aversive stimuli but also display different dynamics across learning. NAc Core dopamine signals tracked with successful avoidance learning and were consistent with the encoding of a safety prediction error.^16^ NAc vmShell dopamine signals faded with learning and were consistent with the concept of salience encoding.^12^ We also found that NAc Core dopamine axon activity does not depend on molecular identity and diverges from extracellular dopamine dynamics at high-performance levels. These results suggest that post-release dopamine clearance mechanisms may contribute to avoidance learning-related plasticity in NAc Core dopamine circuits.

Our data agree with previous work reporting that Core dopamine decreases and vmShell dopamine increases for shocks and shock-predicting cues,^7,21,49,50^ confirming that this heterogeneity in response to aversive stimuli relates strongly to anatomy. However, given recent interest in how molecular markers may further define dopamine neuron functional computations,^46^ we were surprised to find that the activity of Core-projecting dopamine axons was consistent regardless of Sox6 expression (Figure 8). We conclude that Sox6 likely plays a stronger role in developmental targeting events than it does in functional properties in mature dopamine neurons.^34^ Given that both the Sox6+ and -populations may be further molecularly subdivided, it is conceivable that other molecular markers might define important subpopulations relevant for learning from aversive outcomes.

Given that we did not find a strong molecular marker for positive vs. negative-going dopamine neuron responses to aversive stimuli, more research is required to determine the genes and resulting physiological or anatomical differences amongst dopamine neuron subpopulations that contribute to subregional differences in dopamine release. Although numerous studies have detected positive-going dopamine signals in response to aversive events,^5–8^ debate continues about the specific locations where these responses occur. In our experiments, Core and vmShell dopamine signals are vastly different, but efforts to identify variables contributing to discrepancies across labs (e.g., subtle differences in mouse vs rat NAc subregion boundaries) would help the field coalesce. Agreement in defining anatomical subregions could be aided by the discovery of reliable molecular markers for projection-targeted dopamine neurons that are directly related to physiological function.

Our data illustrate not only how Core and vmShell dopamine signal vary by subregion, but how their computations relate to learned behavioral strategies as mice transition from naïve to expert avoidance. This perspective allows us to move beyond arguments regarding valence to consider how dopamine computations coordinate across subtypes to promote continuously evolving responses to new situations.

NAc Core dopamine is thought to be crucial for learning cue-outcome associations during reward learning tasks.^28–30^ Our data indicate that instrumental cue-response associations are also encoded during avoidance learning. Evaluating this encoding over days of learning, we found that NAc Core signals relate to avoidance behavior across learning, with improving representations of the warning cue predicting better avoidance of the aversive outcome (Figure 4). These results, in combination with evidence from the prior literature,^16,51^ lead us to conclude that dopamine release in the NAc Core is reflecting learning and consolidation of avoidance rules via safety prediction error encoding.

NAc Core dopamine representations of the warning cue are negative-going. This suppression of dopamine release during the warning cue could be important for downstream synaptic plasticity. Low dopamine causes long-term potentiation of indirect pathway corticostriatal synapses.^11,52–54^ When we recorded calcium responses from NAc Core D1-and D2-SPNs, we found largely similar activity patterns in these two cell types, as has been reported previously.^55,56^ However, D1-SPNs showed the largest responses early in training, whereas D2-SPNs showed their largest responses later (Figure S4). Although these changes in fluorescence could be due to non-somatic changes in calcium,^57^ our findings are consistent with recent work on the NAc microcircuits required for aversive learning, which showed that D1-SPN activity promotes later potentiation at glutamatergic synapses onto D2-SPNs through interactions with cholinergic interneurons.^58,59^

Our finding that vmShell dopamine increases in response to aversive stimuli is consistent with prior research showing positive-going dopamine responses to both rewarding and aversive stimuli in this region.^7,21,50^ However, we found that these positive-going responses were not sustained in the avoidance learning task. One plausible explanation for these findings could be salience. On Day 1, shocks are unpredicted and highly salient, but that saliency would quickly dissipate and transfer to the warning cue as cue-outcome associations were learned on Days 2 and 3. Salience is an important proposed function of positive-going aversive-responsive dopamine signals in the NAc, with other strong experimental and computation evidence.^11,12^ Such salience signals can support learning by prompting attention but may not necessarily be required for rule learning, especially with regard to instrumental behavior. Indeed, our modeling suggests that the overall representation of task events by vmShell dopamine is almost completely gone by Day 7, when rules are well-learned by nearly all mice (Figure 4).

We chose to study avoidance learning not only because it is an instrumental behavior, but also because of its relationship to stress-related psychiatric disorders. Although aversive stimuli are stressful, the harms of stress to mental health are amplified by the loss of a sense of control.^60,61^ The ability to learn avoidance rules can provide a sense of control, whereas poor avoidance learning could promote learned helplessness and depression.^62,63^ Conversely, inappropriately strong or overgeneralized avoidance learning may be related to anxiety disorders and OCD.^2,64,65^ To better understand how the dopamine signals we recorded during controllable aversive stimuli would compare to a situation where control is removed, at the end of avoidance training, we submitted mice to a day of inescapable shocks. We found that the controllability of the stressor changes dopamine responses in both Core and vmShell. In Core, the magnitude of responses to the cue decreased while in vmShell the magnitude increased. Additionally, very strong phasic release of dopamine was evoked by the cessation of the inescapable shock in both regions. Previous research has shown that VTA dopamine neuron responses at punishment termination are important for motivated behavior to escape punishment and can be diminished by learned helplessness paradigms.^66^ However, regional variation of dopamine signals at shock termination, and the information encoded by such signals for use by downstream circuits, is not clear. Although the large dopamine signals we observed in response to inescapable shock termination in Core and vmShell may result from similar mechanisms or extracellular overflow between regions, they could also represent fundamentally different information for each region – that is, these signals are consistent both with unpredicted safety in the Core and salience in the vmShell. Further research to understand these dopamine signals, how they are generated (perhaps in part by differing inputs from limbic centers such as the amygdala, ventral hippocampus, and hypothalamus,^21,22,67,68^ and how they might vary in rodent models of anxiety, depression, and OCD would be revealing.

In conclusion, our work supports a model for NAc Core and vmShell dopamine function in which these two regions not only encode aversion oppositely but also guide avoidance through different computational principles. Further understanding this computational heterogeneity in dopamine function, and its behavioral consequences, can provide avenues to understand how dopamine subpopulations work in concert to provide animals with tools to avoid danger in their environment.

## Acknowledgments

We thank the members of the Lerner laboratory for helpful discussions and critical feedback throughout the project. We thank the Center for Comparative Medicine at Northwestern University for providing care for all mice used in these studies. We thank the members of the Center for Translational Pain Research, particularly Drs. Vania Apkarian, Marco Martina, and Jones Parker, for input, feedback, and support on these studies.

## Author Contributions

G.C.L. and T.N.L. conceived the experiments, and G.C.L., L.D.V.C., and O.A.M-R. executed them. G.C.L., L.D.V.C., R.F.K., M.D.S., J.C., and T.N.L. analyzed the data, including modeling studies. R.A., J.C., and T.N.L. provided oversight and support for the project. G.C.L. and T.N.L. wrote the manuscript with assistance from the other authors.

## Supplemental information

Document S1. Figures S1–S5 and Tables S1-S3

## References

1. Hofmann, S.G., and Hay, A.C. (2018). Rethinking Avoidance: Toward a Balanced Approach to Avoidance in Treating Anxiety Disorders. J Anxiety Disord 55, 14–21. 10.1016/j.janxdis.2018.03.004.

2. Ball, T.M., and Gunaydin, L.A. (2022). Measuring maladaptive avoidance: from animal models to clinical anxiety. Neuropsychopharmacol. 47, 978–986. 10.1038/s41386-021-01263-4.

3. Wise, R.A. (2004). Dopamine, learning and motivation. Nat Rev Neurosci 5, 483–494. 10.1038/nrn1406.

4. Schultz, W. (1997). Dopamine neurons and their role in reward mechanisms. Current Opinion in Neurobiology 7, 191–197. 10.1016/S0959-4388(97)80007-4.

5. Matsumoto, M., and Hikosaka, O. (2009). Two types of dopamine neuron distinctly convey positive and negative motivational signals. Nature 459, 837–841. 10.1038/nature08028.

6. Brischoux, F., Chakraborty, S., Brierley, D.I., and Ungless, M.A. (2009). Phasic excitation of dopamine neurons in ventral VTA by noxious stimuli. Proceedings of the National Academy of Sciences 106, 4894–4899. 10.1073/pnas.0811507106.

7. Yuan, L., Dou, Y.-N., and Sun, Y.-G. (2019). Topography of Reward and Aversion Encoding in the Mesolimbic Dopaminergic System. J. Neurosci. 39, 6472–6481. 10.1523/JNEUROSCI.0271-19.2019.

8. Lerner, T.N., Shilyansky, C., Davidson, T.J., Evans, K.E., Beier, K.T., Zalocusky, K.A., Crow, A.K., Malenka, R.C., Luo, L., Tomer, R., et al. (2015). Intact-Brain Analyses Reveal Distinct Information Carried by SNc Dopamine Subcircuits. Cell 162, 635–647. 10.1016/j.cell.2015.07.014.

9. Verharen, J.P.H., Zhu, Y., and Lammel, S. (2020). Aversion hot spots in the dopamine system. Current Opinion in Neurobiology 64, 46–52. 10.1016/j.conb.2020.02.002.

10. Saddoris, M.P., Cacciapaglia, F., Wightman, R.M., and Carelli, R.M. (2015). Differential Dopamine Release Dynamics in the Nucleus Accumbens Core and Shell Reveal Complementary Signals for Error Prediction and Incentive Motivation. J Neurosci 35, 11572–11582. 10.1523/JNEUROSCI.2344-15.2015.

11. Bromberg-Martin, E.S., Matsumoto, M., and Hikosaka, O. (2010). Dopamine in Motivational Control: Rewarding, Aversive, and Alerting. Neuron 68, 815–834. 10.1016/j.neuron.2010.11.022.

12. Kutlu, M.G., Zachry, J.E., Melugin, P.R., Cajigas, S.A., Chevee, M.F., Kelly, S.J., Kutlu, B., Tian, L., Siciliano, C.A., and Calipari, E.S. (2021). Dopamine release in the nucleus accumbens core signals perceived saliency. Current Biology 31, 4748–4761.e8. 10.1016/j.cub.2021.08.052.

13. Watabe-Uchida, M., Eshel, N., and Uchida, N. (2017). Neural Circuitry of Reward Prediction Error. Annual Review of Neuroscience 40, 373–394. 10.1146/annurev-neuro-072116-031109.

14. Lerner, T.N., Holloway, A.L., and Seiler, J.L. (2021). Dopamine, Updated: Reward Prediction Error and Beyond. Current Opinion in Neurobiology 67, 123–130. 10.1016/j.conb.2020.10.012.

15. Oleson, E.B., Gentry, R.N., Chioma, V.C., and Cheer, J.F. (2012). Subsecond Dopamine Release in the Nucleus Accumbens Predicts Conditioned Punishment and Its Successful Avoidance. J. Neurosci. 32, 14804–14808. 10.1523/JNEUROSCI.3087-12.2012.

16. Stelly, C.E., Haug, G.C., Fonzi, K.M., Garcia, M.A., Tritley, S.C., Magnon, A.P., Ramos, M.A.P., and Wanat, M.J. (2019). Pattern of dopamine signaling during aversive events predicts active avoidance learning. Proc Natl Acad Sci U S A 116, 13641–13650. 10.1073/pnas.1904249116.

17. Chou, S.-H., Chen, Y.-J., Liao, C.-P., and Pan, C.-L. (2022). A role for dopamine in *C. elegans* avoidance behavior induced by mitochondrial stress. Neuroscience Research 178, 87–92. 10.1016/j.neures.2022.01.005.

18. Akiti, K., Tsutsui-Kimura, I., Xie, Y., Mathis, A., Markowitz, J.E., Anyoha, R., Datta, S.R., Mathis, M.W., Uchida, N., and Watabe-Uchida, M. (2022). Striatal dopamine explains novelty-induced behavioral dynamics and individual variability in threat prediction. Neuron 110, 3789–3804.e9. 10.1016/j.neuron.2022.08.022.

19. Wenzel, J.M., Oleson, E.B., Gove, W.N., Cole, A.B., Gyawali, U., Dantrassy, H.M., Bluett, R.J., Dryanovski, D.I., Stuber, G.D., Deisseroth, K., et al. (2018). Phasic Dopamine Signals in the Nucleus Accumbens that Cause Active Avoidance Require Endocannabinoid Mobilization in the Midbrain. Current Biology 28, 1392–1404.e5. 10.1016/j.cub.2018.03.037.

20. Hollon, N.G., Soden, M.E., and Wanat, M.J. (2013). Dopaminergic Prediction Errors Persevere in the Nucleus Accumbens Core during Negative Reinforcement. J. Neurosci. 33, 3253–3255. 10.1523/JNEUROSCI.5762-12.2013.

21. 21. de Jong, J.W., Afjei, S.A., Pollak Dorocic, I., Peck, J.R., Liu, C., Kim, C.K., Tian, L., Deisseroth, K., and Lammel, S. (2019). A Neural Circuit Mechanism for Encoding Aversive Stimuli in the Mesolimbic Dopamine System. Neuron 101, 133–151.e7. 10.1016/j.neuron.2018.11.005.

22. Li, Z., Chen, Z., Fan, G., Li, A., Yuan, J., and Xu, T. (2018). Cell-Type-Specific Afferent Innervation of the Nucleus Accumbens Core and Shell. Front Neuroanat 12, 84. 10.3389/fnana.2018.00084.

23. Castro, D.C., and Bruchas, M.R. (2019). A Motivational and Neuropeptidergic Hub: Anatomical and Functional Diversity within Nucleus Accumbens Shell. Neuron 102, 529–552. 10.1016/j.neuron.2019.03.003.

24. Peleg-Raibstein, D., and Feldon, J. (2006). Effects of dorsal and ventral hippocampal NMDA stimulation on nucleus accumbens core and shell dopamine release. Neuropharmacology 51, 947–957. 10.1016/j.neuropharm.2006.06.002.

25. French, S.J., and Totterdell, S. (2003). Individual nucleus accumbens-projection neurons receive both basolateral amygdala and ventral subicular afferents in rats. Neuroscience 119, 19–31. 10.1016/S0306-4522(03)00150-7.

26. Ikemoto, S. (2007). Dopamine reward circuitry: Two projection systems from the ventral midbrain to the nucleus accumbens–olfactory tubercle complex. Brain Research Reviews 56, 27–78. 10.1016/j.brainresrev.2007.05.004.

27. Collins, A.L., and Saunders, B.T. (2020). Heterogeneity in striatal dopamine circuits: Form and function in dynamic reward seeking. Journal of Neuroscience Research 98, 1046–1069. 10.1002/jnr.24587.

28. Ambroggi, F., Ghazizadeh, A., Nicola, S.M., and Fields, H.L. (2011). Roles of Nucleus Accumbens Core and Shell in Incentive-Cue Responding and Behavioral Inhibition. J Neurosci 31, 6820–6830. 10.1523/JNEUROSCI.6491-10.2011.

29. Saddoris, M.P. (2013). Rapid dopamine dynamics in the accumbens core and shell Learning and action. Front Biosci E5, 273–288. 10.2741/E615.

30. Floresco, S.B. (2015). The Nucleus Accumbens: An Interface Between Cognition, Emotion, and Action. Annual Review of Psychology 66, 25–52. 10.1146/annurev-psych-010213-115159.

31. Taira, M., Millard, S.J., Verghese, A., DiFazio, L.E., Hoang, I.B., Jia, R., Sias, A., Wikenheiser, A., and Sharpe, M.J. (2024). Dopamine Release in the Nucleus Accumbens Core Encodes the General Excitatory Components of Learning. J. Neurosci. 10.1523/JNEUROSCI.0120-24.2024.

32. Day, J.J., Roitman, M.F., Wightman, R.M., and Carelli, R.M. (2007). Associative learning mediates dynamic shifts in dopamine signaling in the nucleus accumbens. Nature Neuroscience 10, 1020–1028. 10.1038/nn1923.

33. Poulin, J.-F., Caronia, G., Hofer, C., Cui, Q., Helm, B., Ramakrishnan, C., Chan, C.S., Dombeck, D., Deisseroth, K., and Awatramani, R. (2018). Mapping projections of molecularly defined dopamine neuron subtypes using intersectional genetic approaches. Nat Neurosci 21, 1260–1271. 10.1038/s41593-018-0203-4.

34. Pereira Luppi, M., Azcorra, M., Caronia-Brown, G., Poulin, J.-F., Gaertner, Z., Gatica, S., Moreno-Ramos, O.A., Nouri, N., Dubois, M., Ma, Y.C., et al. (2021). Sox6 expression distinguishes dorsally and ventrally biased dopamine neurons in the substantia nigra with distinctive properties and embryonic origins. Cell Reports 37, 109975. 10.1016/j.celrep.2021.109975.

35. Pereira, T.D., Tabris, N., Matsliah, A., Turner, D.M., Li, J., Ravindranath, S., Papadoyannis, E.S., Normand, E., Deutsch, D.S., Wang, Z.Y., et al. (2022). SLEAP: A deep learning system for multi-animal pose tracking. Nat Methods 19, 486–495. 10.1038/s41592-022-01426-1.

36. Goodwin, N.L., Choong, J.J., Hwang, S., Pitts, K., Bloom, L., Islam, A., Zhang, Y.Y., Szelenyi, E.R., Tong, X., Newman, E.L., et al. (2024). Simple Behavioral Analysis (SimBA) as a platform for explainable machine learning in behavioral neuroscience. Nat Neurosci 27, 1411–1424. 10.1038/s41593-024-01649-9.

37. Sherathiya, V.N., Schaid, M.D., Seiler, J.L., Lopez, G.C., and Lerner, T.N. (2021). GuPPy, a Python toolbox for the analysis of fiber photometry data. Sci Rep 11, 24212. 10.1038/s41598-021-03626-9.

38. Ramsay, J.O. Matlab, R and S-PLUS Functions for Functional Data Analysis.

39. Karim, F., Majumdar, S., Darabi, H., and Harford, S. (2019). Multivariate LSTM-FCNs for Time Series Classification. Neural Networks 116, 237–245. 10.1016/j.neunet.2019.04.014.

40. Wu, J., Chen, X.-Y., Zhang, H., Xiong, L.-D., Lei, H., and Deng, S.-H. (2019). Hyperparameter Optimization for Machine Learning Models Based on Bayesian Optimizationb. Journal of Electronic Science and Technology 17, 26–40. 10.11989/JEST.1674-862X.80904120.

41. Comet.ml Comet.ml -Supercharging Machine Learning. Comet.ml - Supercharging Machine Learning. https://www.comet.com/mschaid/lstmnfcn-bayes-tuning-5-day-weighted/view/new/panels.

42. Cox, J., Minerva, A.R., Fleming, W.T., Zimmerman, C.A., Hayes, C., Zorowitz, S., Bandi, A., Ornelas, S., McMannon, B., Parker, N.F., et al. (2023). A neural substrate of sex-dependent modulation of motivation. Nat Neurosci 26, 274–284. 10.1038/s41593-022-01229-9.

43. Engelhard, B., Finkelstein, J., Cox, J., Fleming, W., Jang, H.J., Ornelas, S., Koay, S.A., Thiberge, S.Y., Daw, N.D., Tank, D.W., et al. (2019). Specialized coding of sensory, motor and cognitive variables in VTA dopamine neurons. Nature 570, 509–513. 10.1038/s41586-019-1261-9.

44. Choi, J.Y., Jang, H.J., Ornelas, S., Fleming, W.T., Fürth, D., Au, J., Bandi, A., Engel, E.A., and Witten, I.B. (2020). A Comparison of Dopaminergic and Cholinergic Populations Reveals Unique Contributions of VTA Dopamine Neurons to Short-Term Memory. Cell Reports 33. 10.1016/j.celrep.2020.108492.

45. Pinto, L., and Dan, Y. (2015). Cell-Type-Specific Activity in Prefrontal Cortex during Goal-Directed Behavior. Neuron 87, 437–450. 10.1016/j.neuron.2015.06.021.

46. Azcorra, M., Gaertner, Z., Davidson, C., He, Q., Kim, H., Nagappan, S., Hayes, C.K., Ramakrishnan, C., Fenno, L., Kim, Y.S., et al. (2023). Unique functional responses differentially map onto genetic subtypes of dopamine neurons. Nat Neurosci 26, 1762– 1774. 10.1038/s41593-023-01401-9.

47. Mohebi, A., Pettibone, J.R., Hamid, A.A., Wong, J.-M.T., Vinson, L.T., Patriarchi, T., Tian, L., Kennedy, R.T., and Berke, J.D. (2019). Dissociable dopamine dynamics for learning and motivation. Nature 570, 65–70. 10.1038/s41586-019-1235-y.

48. Liu, C., Cai, X., Ritzau-Jost, A., Kramer, P.F., Li, Y., Khaliq, Z.M., Hallermann, S., and Kaeser, P.S. (2022). An action potential initiation mechanism in distal axons for the control of dopamine release. Science 375, 1378–1385. 10.1126/science.abn0532.

49. Salinas-Hernández, X.I., Zafiri, D., Sigurdsson, T., and Duvarci, S. (2023). Functional architecture of dopamine neurons driving fear extinction learning. Neuron 111, 3854–3870.e5. 10.1016/j.neuron.2023.08.025.

50. Badrinarayan, A., Wescott, S.A., Weele, C.M.V., Saunders, B.T., Couturier, B.E., Maren, S., and Aragona, B.J. (2012). Aversive Stimuli Differentially Modulate Real-Time Dopamine Transmission Dynamics within the Nucleus Accumbens Core and Shell. J. Neurosci. 32, 15779–15790. 10.1523/JNEUROSCI.3557-12.2012.

51. Ray, M.H., Russ, A.N., Walker, R.A., and McDannald, M.A. (2020). The Nucleus Accumbens Core is Necessary to Scale Fear to Degree of Threat. J. Neurosci. 40, 4750– 4760. 10.1523/JNEUROSCI.0299-20.2020.

52. Lerner, T.N., and Kreitzer, A.C. (2011). Neuromodulatory control of striatal plasticity and behavior. Current Opinion in Neurobiology 21, 322–327. 10.1016/j.conb.2011.01.005.

53. Surmeier, D.J., Carrillo-Reid, L., and Bargas, J. (2011). Dopaminergic modulation of striatal neurons, circuits, and assemblies. Neuroscience 198, 3–18. 10.1016/j.neuroscience.2011.08.051.

54. Shen, W., Flajolet, M., Greengard, P., and Surmeier, D.J. (2008). Dichotomous dopaminergic control of striatal synaptic plasticity. Science 321, 848–851. 10.1126/science.1160575.

55. Tecuapetla, F., Jin, X., Lima, S.Q., and Costa, R.M. (2016). Complementary Contributions of Striatal Projection Pathways to Action Initiation and Execution. Cell 166, 703–715. 10.1016/j.cell.2016.06.032.

56. Parker, J.G., Marshall, J.D., Ahanonu, B., Wu, Y.-W., Kim, T.H., Grewe, B.F., Zhang, Y., Li, J.Z., Ding, J.B., Ehlers, M.D., et al. (2018). Diametric neural ensemble dynamics in parkinsonian and dyskinetic states. Nature 557, 177–182. 10.1038/s41586-018-0090-6.

57. Legaria, A.A., Matikainen-Ankney, B.A., Yang, B., Ahanonu, B., Licholai, J.A., Parker, J.G., and Kravitz, A.V. (2022). Fiber photometry in striatum reflects primarily nonsomatic changes in calcium. Nat Neurosci 25, 1124–1128. 10.1038/s41593-022-01152-z.

58. Belilos, A., Gray, C., Sanders, C., Black, D., Mays, E., Richie, C., Sengupta, A., Hake, H., and Francis, T.C. (2023). Nucleus accumbens local circuit for cue-dependent aversive learning. Cell Reports 42, 113488. 10.1016/j.celrep.2023.113488.

59. Francis, T.C., Yano, H., Demarest, T.G., Shen, H., and Bonci, A. (2019). High-Frequency Activation of Nucleus Accumbens D1-MSNs Drives Excitatory Potentiation on D2-MSNs. Neuron 103, 432–444.e3. 10.1016/j.neuron.2019.05.031.

60. Riachi, E., Holma, J., and Laitila, A. (2024). Psychotherapists’ perspectives on loss of sense of control. Brain and Behavior 14. 10.1002/brb3.3368.

61. Hancock, L., and Bryant, R.A. (2020). Posttraumatic stress, stressor controllability, and avoidance. Behaviour Research and Therapy 128, 103591. 10.1016/j.brat.2020.103591.

62. Chase, H.W., Frank, M.J., Michael, A., Bullmore, E.T., Sahakian, B.J., and Robbins, T.W. (2010). Approach and avoidance learning in patients with major depression and healthy controls: relation to anhedonia. Psychological Medicine 40, 433–440. 10.1017/S0033291709990468.

63. Grant, D.M., Wingate, L.R., Rasmussen, K.A., Davidson, C.L., Slish, M.L., Rhoades-Kerswill, S., Mills, A.C., and Judah, M.R. (2013). An Examination of the Reciprocal Relationship Between Avoidance Coping and Symptoms of Anxiety and Depression. Journal of Social and Clinical Psychology 32, 878–896. 10.1521/jscp.2013.32.8.878.

64. Endrass, T., Kloft, L., Kaufmann, C., and Kathmann, N. (2011). Approach and avoidance learning in obsessive–compulsive disorder. Depression and Anxiety 28, 166–172. 10.1002/da.20772.

65. Geramita, M.A., Yttri, E.A., and Ahmari, S.E. (2020). The two-step task, avoidance, and OCD. Journal of Neuroscience Research 98, 1007–1019. 10.1002/jnr.24594.

66. Wu, M., Minkowicz, S., Dumrongprechachan, V., Hamilton, P., Xiao, L., and Kozorovitskiy, Y. (2021). Attenuated dopamine signaling after aversive learning is restored by ketamine to rescue escape actions. eLife 10, e64041. 10.7554/eLife.64041.

67. Watabe-Uchida, M., Zhu, L., Ogawa, S.K., Vamanrao, A., and Uchida, N. (2012). Whole-Brain Mapping of Direct Inputs to Midbrain Dopamine Neurons. Neuron 74, 858–873. 10.1016/j.neuron.2012.03.017.

68. Beier, K.T., Steinberg, E.E., DeLoach, K.E., Xie, S., Miyamichi, K., Schwarz, L., Gao, X.J., Kremer, E.J., Malenka, R.C., and Luo, L. (2015). Circuit Architecture of VTA Dopamine Neurons Revealed by Systematic Input-Output Mapping. Cell 162, 622–634. 10.1016/j.cell.2015.07.015.

