## Supplemental Information for "Region-specific Nucleus Accumbens Dopamine Signals Encode Distinct Aspects of Avoidance Learning"

Figure S1

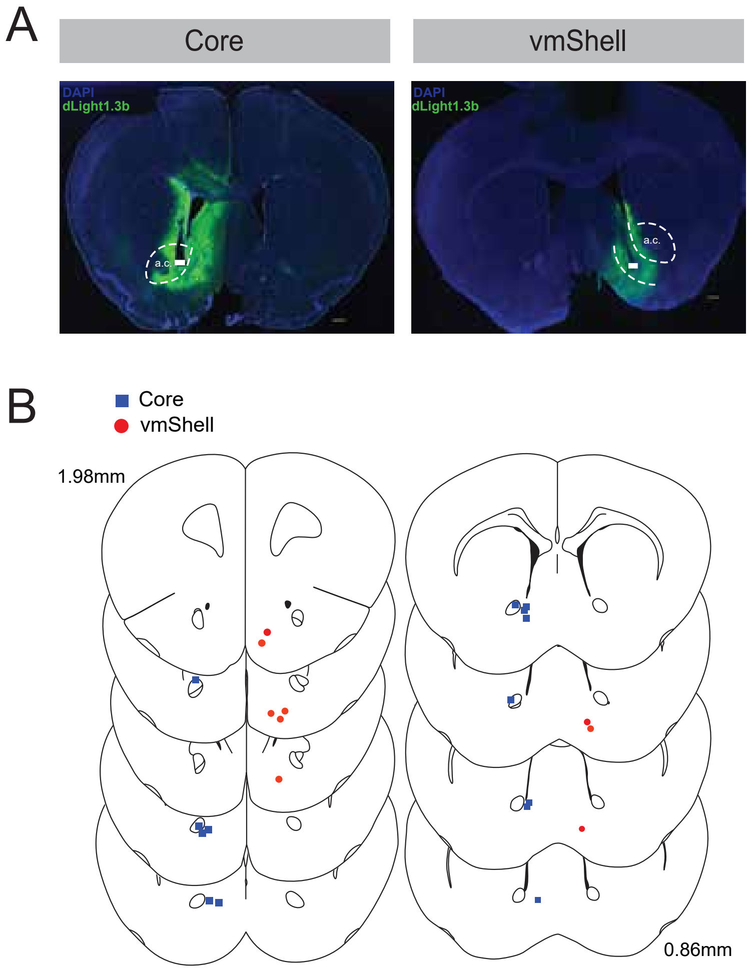
A
Core
vmShell
DAPI
dLight1.3b
a.c.
DAPI
dLight1.3b
a.c.
B
Core
vmShell
1.98mm
0.86mm

**S1. Recording sites for dLight1.3b recordings, related to Figures 2-7**

(A) Representative histology images (4x) indicating viral spread of dLight 1.3b (green, all images) and probe placement for Core (left) and vmShell (right). DAPI (blue, all images) was used for nuclear staining. Scale bars are 300 µm; a.c., anterior commissure. Thick white bars indicate probe placement.

(B) Probe placements in Core (blue squares) and vmShell (red circles) for all mice included in Figures 1-7.

Figure S2

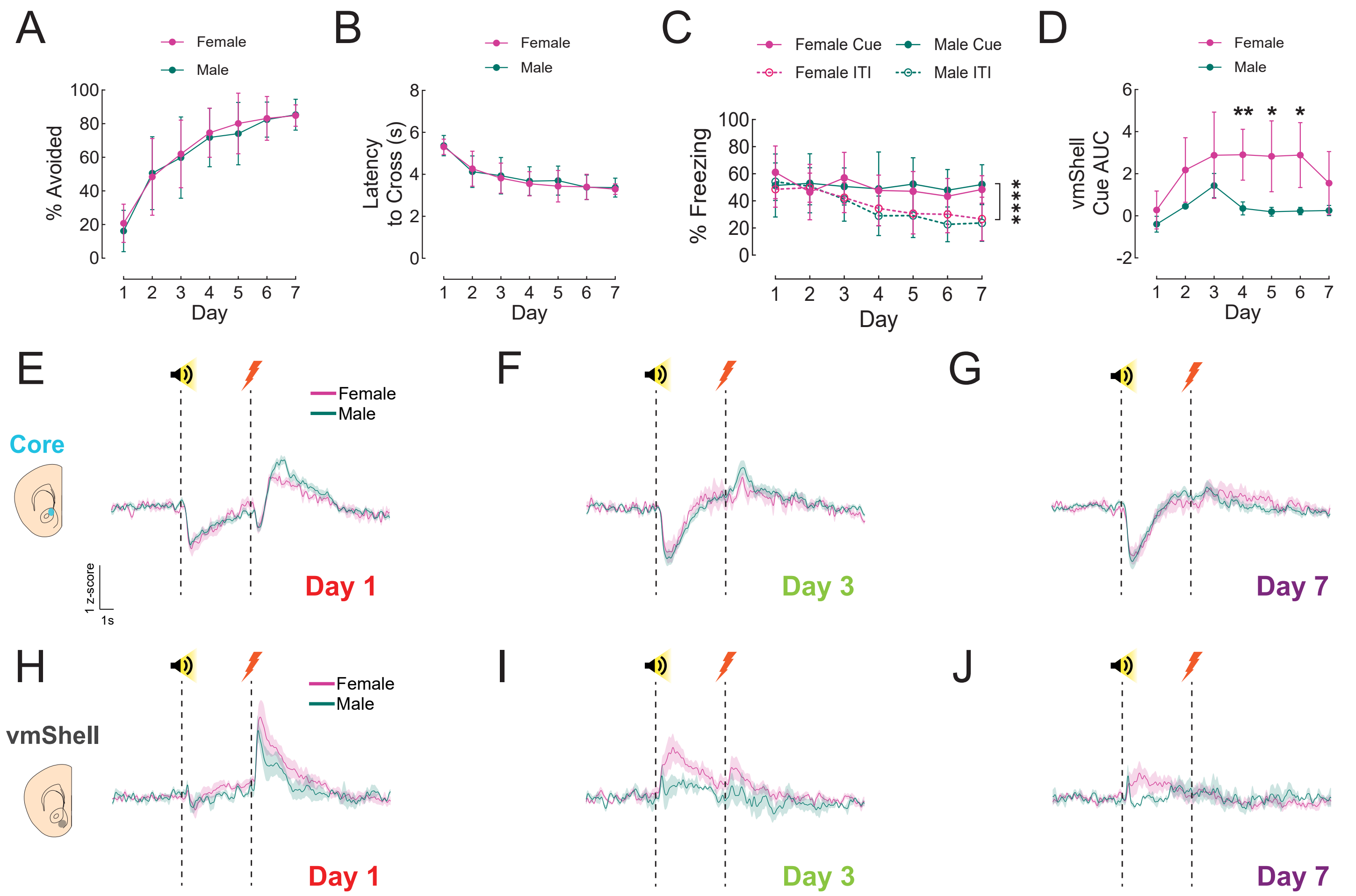

A
Female
Male
% Avoided
Day
B
Latency to Cross (s)
Day
C
Female Cue
Male Cue
Female ITI
Male ITI
% Freezing
Day
****
D
vmShell Cue AUC
**
*
Day
E
Female
Male
Core
1 z-score
1s
Day 1
F
Day 3
G
Day 7
H
vmShell
Day 1
I
Day 3
J
Day 7

**S2. Lack of sex differences in NAc Core and vmShell dopamine responses during avoidance learning, related to Figures 1-2**

(A) Percentage of shocks avoided over days of active avoidance training split by sex. No significant differences are seen between females (n = 18, pink) and males (n = 18, green) across days of training.

(B) Latency to cross over days of active avoidance training split by sex. No significant differences are seen between females (n = 18, pink) and males (n = 18, green) across days of training.

(C) Percentage of freezing during the cue (solid line) and ITI (dashed line) over days of active avoidance split by sex. While there is a significant difference for freezing between cue and ITI (****$p<0.0001$), no significant differences are seen between females (n = 12, pink) and males (n = 13, green) across days of training.

(D) Cue AUC for vmShell dLight1.3b signals over days split by sex (females n = 5, males n = 3). Days 4, 5, and 6 are significantly different by sex. *$p<0.05$, **$p<0.01$.

(E-G) Plots showing the dopamine signals collected from Core for females (n = 5, pink) and males (n = 9, green) aligned to the start of the warning cue (first dotted line). Shock occurred 5s after the warning cue start on escape trials (second dotted line). Day 1 (E), Day 3 (F), and Day 7 (G) are shown.

(H-J) Plots showing the dopamine signals collected from vmShell for females (n = 5, pink) and males (n = 3, green) aligned to the start of the warning cue (first dotted line). Shock occurred 5s after the warning cue start on escape trials (second dotted line). Day 1 (H), Day 3 (I), and Day 7 (J) are shown.

Figure S3

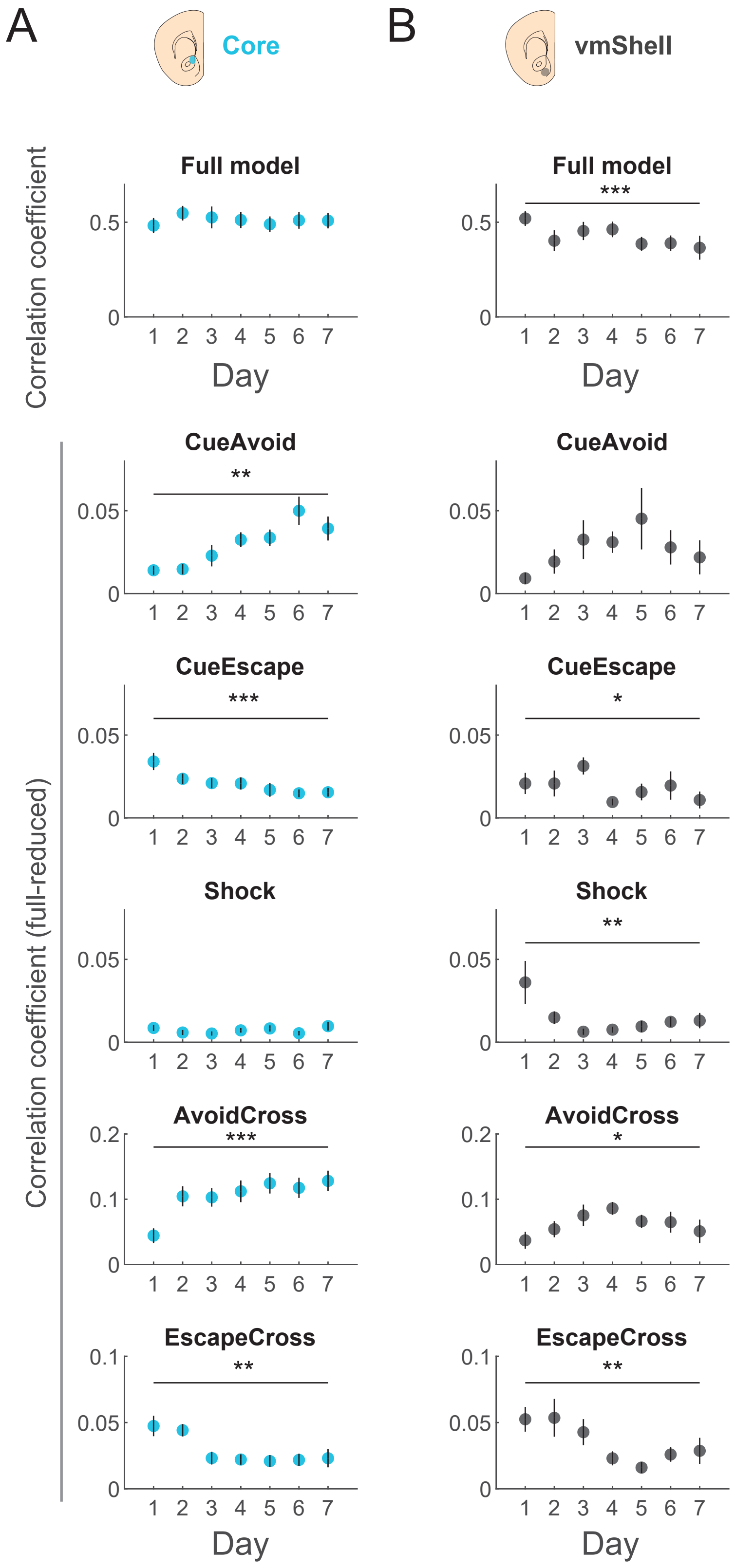
A
Core
B
vmShell
Correlation coefficient
Full model
***
0.5
0
1 2 3 4 5 6 7
Day
Correlation coefficient (full-reduced)
CueAvoid
**
0.05
CueEscape
***
*
Shock
**
AvoidCross
***
*
0.2
0.1
EscapeCross
**
0.1
0.05
Day

**S3. Analysis of encoding of individual events by modeling dopamine fluorescence in individual mice, related to Figure 4**

(A-B *top*) Correlation coefficients for the full encoding model of dopamine fluorescence for Core (A) and vmShell (B) shows a similar conclusion to Figure 4 – that Core encoding remains steady over days, while vmShell encoding weakens.

(A-B *bottom)* The difference in correlation coefficients for the full model compared to a reduced model with each event removed.

*p<0.05, **p<0.01, ***p<0.001 for a change over days by one-way, repeated measures ANOVA. Error bars are standard error of the mean.

Figure S4

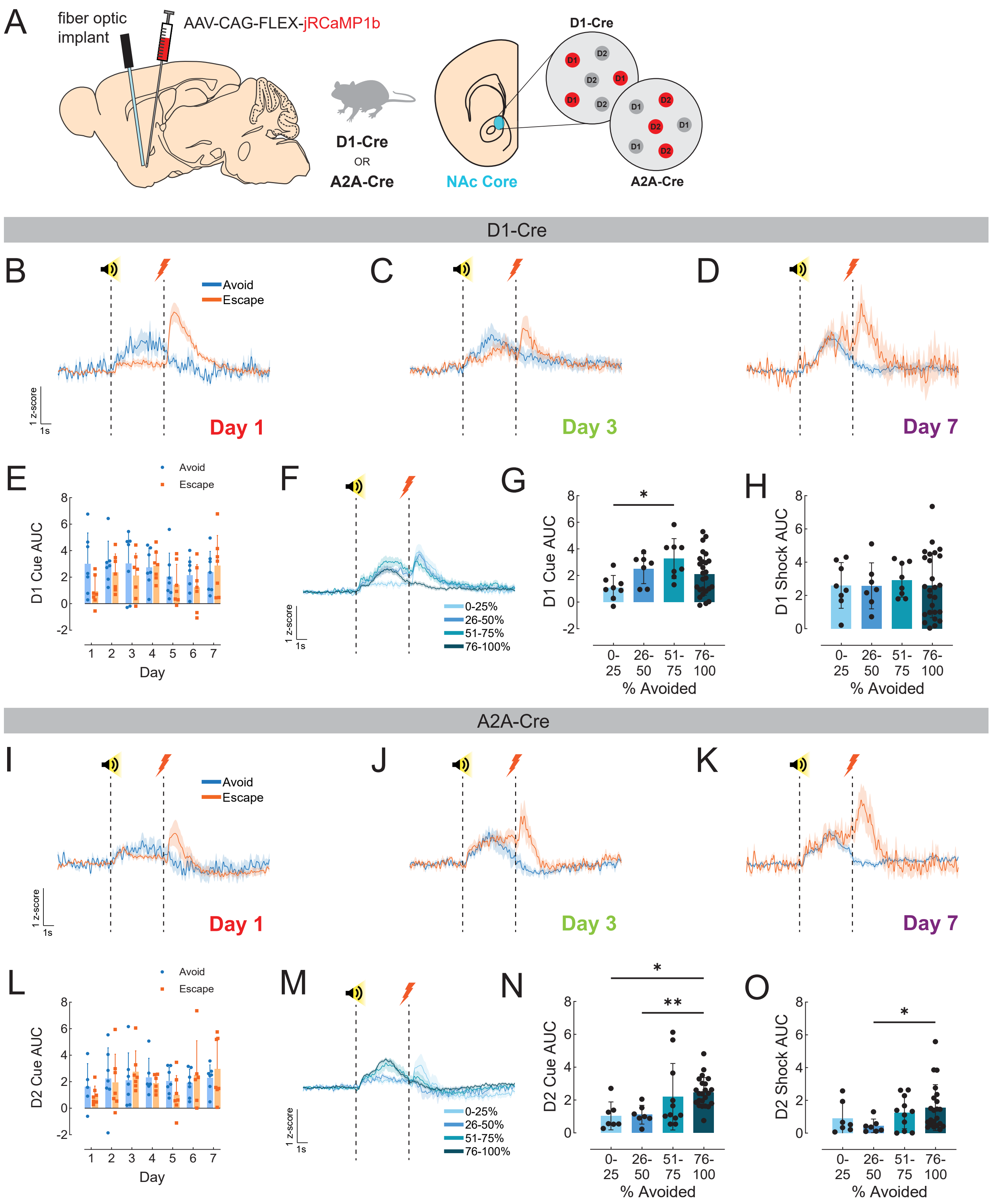

A
fiber optic implant
AAV-CAG-FLEX-jRCaMP1b
D1-Cre
OR
A2A-Cre
NAc Core
D1-Cre
A2A-Cre
D1-Cre
B
C
D
Avoid
Escape
1 z-score
1s
Day 1
Day 3
Day 7
E
Avoid
Escape
D1 Cue AUC
Day
F
0-25%
26-50%
51-75%
76-100%
G
D1 Cue AUC
% Avoided
H
D1 Shock AUC
% Avoided
A2A-Cre
I
J
K
Avoid
Escape
Day 1
Day 3
Day 7
L
D2 Cue AUC
Day
M
0-25%
26-50%
51-75%
76-100%
N
D2 Cue AUC
% Avoided
O
D2 Shock AUC
% Avoided

**S4. Recording from D1- and D2-SPNs in NAc Core during avoidance learning, related to Figures 3 + 6**

(A) Viral strategy for D1- and D2-SPN recordings. An AAV expressing cre-dependent jRCaMP1b was injected into the NAc Core of D1-Cre or A2A-Cre to report calcium activity of D1- or D2-SPNs, respectively. A fiber optic was placed at the same site to allow the collection of fluorescent signals by fiber photometry.

(B-D) Plots showing D1-SPN activity (n = 7) collected during avoid trials (blue) and escape trials (orange) from aligned to the start of the warning cue (first dotted line). Shock occurred 5s after the warning cue start on escape trials (second dotted line). Day 1 (B), Day 3 (C), and Day 7 (D) are shown.

(E) The D1-SPN area-under-the-curve (AUC) for the cue period (0-5s) is shown for avoid and escape trials across days. No significant differences are observed between avoid and escape trials.

(F) Cue-aligned D1-SPN signals divided out according to the avoidance performance of the mouse rather than day recorded.

(G) D1-SPN cue AUC by performance level. *$p<0.05$, 0-25% compared to 51-75% avoidance.

(H) D1-SPN shock AUC by performance level. No significant differences are observed between performance levels.

(I-K) Plots showing D2-SPN activity (n = 7) collected during avoid trials (blue) and escape trials (orange) from aligned to the start of the warning cue (first dotted line). Shock occurred 5s after the warning cue start on escape trials (second dotted line). Day 1 (I), Day 3 (J), and Day 7 (K) are shown.

(L) The D2-SPN area-under-the-curve (AUC) for the cue period (0-5s) is shown for avoid and escape trials across days. No significant differences are observed between avoid and escape trials.

(M) Cue-aligned D2-SPN signals divided out according to the avoidance performance of the mouse rather than day recorded.

(N) D2-SPN cue AUC by performance level. *$p<0.05$, 0-25% compared to 76-100% avoidance, **$p<0.001$, 26-50% compared to 76-100% avoidance.

(O) D2-SPN shock AUC by performance level. *$p<0.05$, 26-50% compared to 76-100% avoidance.

Figure S5

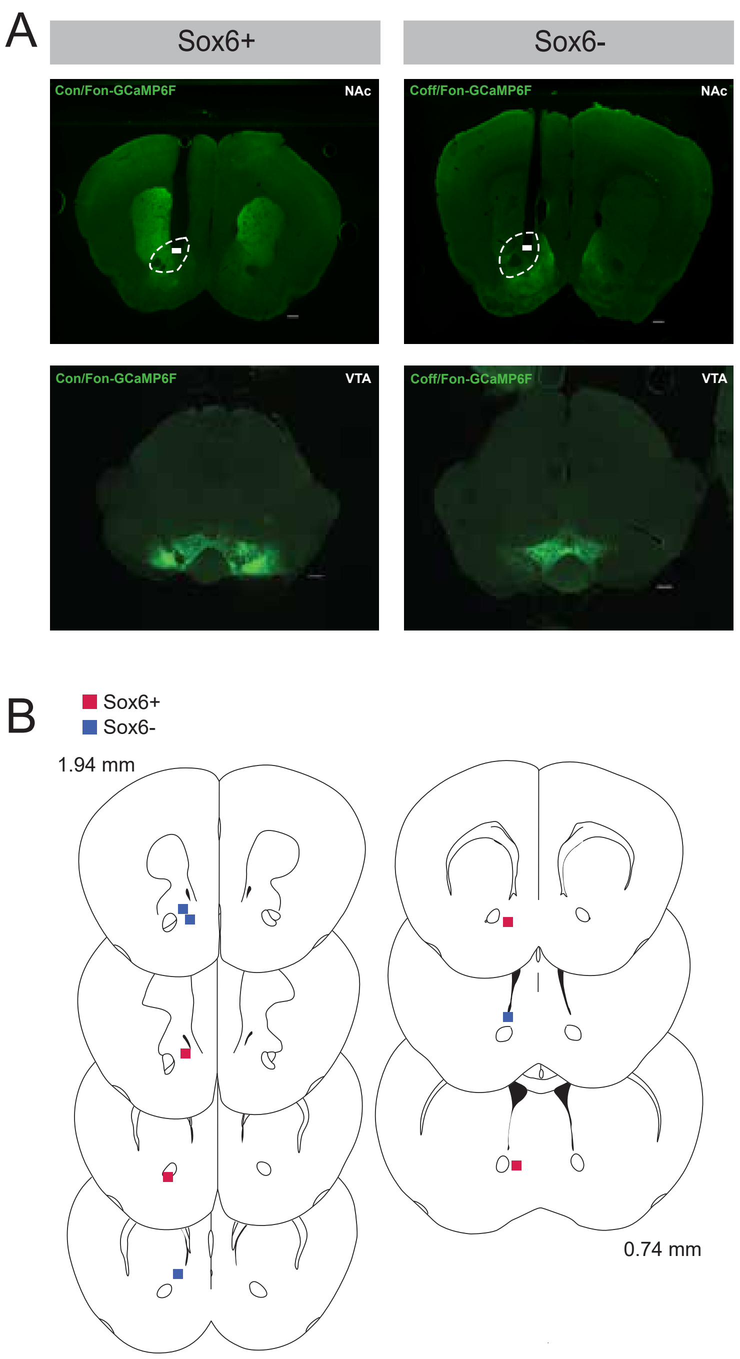
A
Sox6+
Sox6-
Con/Fon-GCaMP6F
NAc
Coff/Fon-GCaMP6F
NAc
Con/Fon-GCaMP6F
VTA
Coff/Fon-GCaMP6F
VTA
B
Sox6+
Sox6-
1.94 mm
0.74 mm

**S5. Recording sites for Sox6+ and Sox6- dopamine axon recordings, related to Figure 8**

(A) Representative histology images (4x) indicating viral spread of Con/Fon-GCaMP6f (green, left images) or Coff/Fon-GCaMP6f (green, right images) and probe placement for Sox6+ (left) or Sox6- (right) expressing dopamine neurons. The top panels are representative images of the NAc while the bottom panels are representative of the VTA. Scale bars are 300 μm. Thick white bars indicate probe placement.

(B) Probe placements in Sox6+ targeted mice (blue squares) and Sox6- targeted mice (red circles) included in Figure 8.

**SUPPLEMENTARY TABLE**

| Parameter | Value |
|---|---|
| adam amsgrad | FALSE |
| adam beta 1 | 0.9 |
| adam beta 2 | 0.999 |
| adam clipnorm | null |
| adam clipvalue | null |
| adam ema momentum | 0.99 |
| adam ema overwrite frequency | null |
| adam epsilon | 1.00E-07 |
| adam global clipnorm | null |
| adam gradient accumulation steps | null |
| adam learning rate | 0.001 |
| adam loss scale factor | null |
| adam use ema | FALSE |
| adam weight decay | null |
| attention | FALSE |
| batch size | 128 |
| callbacks | null |
| categories | auto |
| drop | null |
| dropout | 0.2 |
| dtype | <class 'numpy.float64'> |
| epochs | 2000 |
| feature name combiner | concat |
| filter sizes | [128,256,128] |
| handle unknown | error |
| kernel sizes | [16,10,6] |
| lstm size | 10 |
| max categories | null |
| min frequency | null |
| n epochs | 2000 |
| Optimizer | adam |
| random state | 42 |
| sparse output | FALSE |
| verbose | 0 |

**Table S1.** Parameters of optimized Long Short-Term Memory Fully Convolutional Network Model, related to Figure 4

| | |
|---|---|
| algorithm | bayes |
| maxCombo | 30 |
| objective | maximize |
| metric | test_balanced_accuracy |
| minSampleSize | 700 |
| retryLimit | 20 |
| retryAssignLimit | 0 |
| epochs | 500, 1000, 1500, 2000 |
| dropout | 0.2, 0.4, 0.6, 0.8 |
| kernel_sizes | (32, 15, 9) (16, 10, 6), (8, 5, 3) |
| filter_sizes | (128, 256, 128),(64, 128, 64) |
| lstm_size | 2, 4, 6, 8, 10 |
| random_state | 42 |

**Table S2.** Hyperparameters Search Space Table for Bayesian optimization of LSTM-FCN, Related to Figure 4

| | Dataset | Value |
|---|---|---|
| Accuracy | Test | 0.8742 |
| | Train | 0.9426 |
| Accuracy (weighted) | Test | 0.7273 |
| | Train | 0.8798 |
| F1 Score | Test | 0.5366 |
| | Train | 0.8443 |
| F1 Score (weighted) | Test | 0.8729 |
| | Train | 0.9406 |
| Precision | Test | 0.55 |
| | Train | 0.9276 |
| Precision (weighted) | Test | 0.8717 |
| | Train | 0.942 |
| ROC-AUC | Test | 0.8512 |
| | Train | 0.9821 |
| Recall | Test | 0.5238 |
| | Train | 0.7747 |
| Recall (weighted) | Test | 0.8742 |
| | Train | 0.9426 |

**Table S3.** Performance metrics for LSTM-FCN model, Related to Figure 4
